# A conserved enzyme found in diverse human gut bacteria interferes with anticancer drug efficacy

**DOI:** 10.1101/820084

**Authors:** Peter Spanogiannopoulos, Than S. Kyaw, Ben G. H. Guthrie, Patrick H. Bradley, Joyce V. Lee, Jonathan Melamed, Ysabella Noelle Amora Malig, Kathy N. Lam, Daryll Gempis, Moriah Sandy, Wes Kidder, Erin L. Van Blarigan, Chloe E. Atreya, Alan Venook, Roy R. Gerona, Andrei Goga, Katherine S. Pollard, Peter J. Turnbaugh

## Abstract

Pharmaceuticals are the top predictor of inter-individual variations in gut microbial community structure^1^, consistent with *in vitro* evidence that host-targeted drugs inhibit gut bacterial growth^2^ and are extensively metabolized by the gut microbiome^3,4^. In oncology, bacterial metabolism has been implicated in both drug efficacy^5,6^ and toxicity^7,8^; however, the degree to which bacterial drug sensitivity and metabolism can be driven by conserved pathways also found in mammalian cells remains poorly understood. Here, we show that anticancer fluoropyrimidine drugs inhibit the growth of diverse gut bacterial strains by disrupting pyrimidine metabolism, as in mammalian cells. Select bacteria metabolized 5-fluorouracil (5-FU) to its inactive metabolite dihydrofluorouracil (DHFU), mimicking the major host pathway for drug clearance. The *preTA* operon was necessary and sufficient for 5-FU inactivation in *Escherichia coli*, exhibited high catalytic efficiency for the reductive reaction, decreased the bioavailability and efficacy of oral fluoropyrimidine treatment in mice, and was prevalent in the gut microbiomes of colorectal cancer patients prior to and during treatment. The observed conservation of both the targets and pathways for metabolism of therapeutics across domains highlights the need to distinguish the relative contributions of human and microbial cells to drug disposition^9^, efficacy, and side effect profiles.

---

Despite major innovations in targeted and immunotherapies for cancer, chemotherapeutic drugs like fluoropyrimidines remain the cornerstone of cancer therapy for a variety of cancer types^10^. Fluoropyrimidines have extensive interactions with the human gut microbiome that may have downstream consequences for treatment outcomes. The oral fluoropyrimidine capecitabine (CAP) is extensively metabolized to 5-FU, 5-fluorodeoxyuridine (FUDR), and other metabolites^4^ (**Extended Data Fig. 1a**). CAP meets the FDA criteria for a “highly variable” drug, with a coefficient of variation >30% in intra-subject pharmacokinetic parameters combined with extensive variation between subjects^11^ that cannot be explained by the known host risk factors^12,13^. Adverse reactions to CAP require dose adjustments in ∼35% of patients and complete discontinuation of therapy in ∼10% of patients; GI side effects are common^14,15^. Fluoropyrimidines were designed to target conserved pathways essential for gut bacterial growth^16,17^ and evidence in rats has shown that these drugs alter the gut microbiota^18–20^. Cross-sectional analyses of the human gut microbiome have linked the use of anticancer drugs (including fluoropyrimidines) to changes in gut microbial community structure and function^21–24^. The bacterial metabolism of fluoropyrimidines may also have downstream consequences for host physiology; studies in *Caenorhabditis elegans* have linked genetic differences in their bacterial food source, *E. coli,* to host drug toxicity^6,25,26^. Furthermore, the increasing administration of CAP and other oral fluoropyrimidines^27^ may enhance the potential for interactions with the gut microbiota prior to first pass metabolism and absorption into general circulation^4^.

These observations motivated us to conduct a systematic screen for human gut bacterial sensitivity to three fluoropyrimidine drugs: CAP, 5-FU, and FUDR (**Extended Data Fig. 1a**). We selected 47 sequenced and publicly available bacterial strains, spanning the 6 most abundant phyla found in the human gut (**Supplementary Table 1**). On average, these bacterial species represent 49% of the gut microbiota (**Supplementary Table 1**). Drug sensitivity was evaluated by determining the minimal inhibitory concentration (MIC) in a rich medium that supported the growth of our entire strain collection (**Supplementary Table 1**). There was extensive variation in sensitivity to 5-FU and FUDR (**Fig. 1a** and **Supplementary Table 1**). A total of 13 (5-FU) and 15 (FUDR) bacterial strains tolerated the highest concentration assayed; 12/13 (5-FU) and 12/15 (FUDR) of these strains displayed partial growth inhibition (range: 22.8-96.0% of the growth control; **Supplementary Table 1**). The MICs of 5-FU and FUDR were significantly associated (r=0.475, *p*=0.0008), as expected based on their shared substructure and downstream metabolites (**Extended Data Fig. 1a**). Bacterial phylogeny did not predict sensitivity to 5-FU, with no significant differences between phyla (**Extended Data Fig. 1b**) and multiple nearest neighbors with opposite phenotypes; for example, *Parabacteroides distasonis* (5-FU MIC=300 ng/ml) and *P. merdae* (5-FU MIC>1 mg/ml) (**Fig. 1a**). In contrast, the Bacteroidetes were significantly more sensitive to FUDR relative to either Actinobacteria or Firmicutes (**Extended Data Fig. 1c**), suggesting potential differences in the permeability of these two related fluoropyrimidines. Consistent with a recent report^26^, short-term exposure of *E. coli* and two representative *Bacteroides* (*B. fragilis* and *B. ovatus*) to 5-FU was sufficient to select for drug resistant mutants (MIC>1 mg/ml; **Extended Data Fig. 1d** and **Supplementary Table 2**).

**Figure 1.**
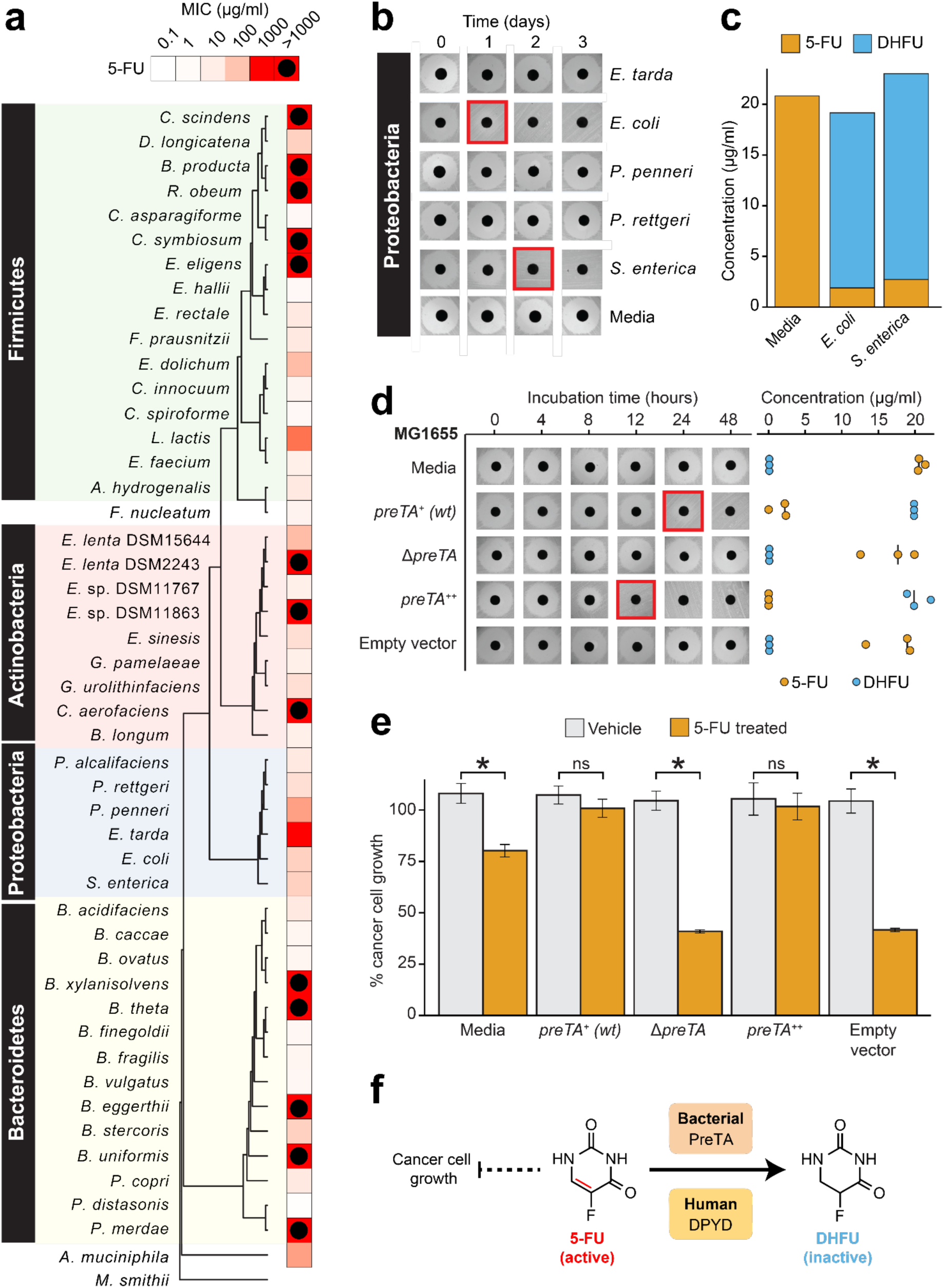
The *preTA* operon is necessary and sufficient for the inactivation of 5-fluorouracil (5-FU) by *E. coli*. **(a)** Heatmap representing the 5-FU minimal inhibitory concentration (MIC, <10% growth relative to vehicle controls; 47 human gut bacterial strains; 2 replicates/strain/concentration; see **Supplementary Table 1**). The phylogenetic tree was built using PhyML^55^, based on a MUSCLE^56^ alignment of full-length 16S rRNA genes. **(b)** We screened 23 strains with MIC≥62.5 µg/ml for drug inactivation using a disk diffusion assay (BHI^+^ media; 20 μg/ml 5-FU); only Proteobacteria are shown. Red squares indicate decreased zones of inhibition. **(c)** LC-QTOF/MS detection of the predicted 5-FU metabolite DHFU after 48 hours of anaerobic incubation [*E. coli* MG1655, *S. enterica* LT2 (DSM17058), 20 μg/ml 5-FU]. **(d)** Wild-type (*preTA^+^*), deletion (*ΔpreTA*), complemented (*preTA^++^*), and empty vector *E. coli* MG1655 strains were assayed for residual 5-FU using disk diffusion (0-48 hours incubation) and LC-QTOF/MS (48 hours). Complementation and empty vector were on the *ΔpreTA* background. Black lines indicate the median value (n=3 biological replicates). **(e)** The *E. coli* MG1655 strains shown in panel d were incubated for 72 hours with 5-FU (20 µg/ml) or vehicle (DMSO, 0.05%), conditioned media was added to the colorectal cell line HCT-116, and cell proliferation was quantified using the MTT assay. Values represent mean±stdev (n=3 biological replicates/strain/condition). \**p*-value<0.001, Student’s *t* test. **(f)** We propose that bacterial PreTA contributes to the elimination of 5-FU similar to hepatic expression of dihydropyrimidine dehydrogenase (DPYD), interfering with drug efficacy.

The bioactivation of the prodrug CAP is thought to be uniquely catalyzed by mammalian enzymes expressed in the intestine and liver^4,28^. Surprisingly, we found that multiple gut bacteria were susceptible to CAP at high but still physiological concentrations (∼14 mg/ml following oral administration)^29^. With the exception of *Providencia rettgeri* we were able to determine a MIC for all of the tested strains (**Supplementary Table 1**). CAP MIC was significantly associated with FUDR MIC (r=0.478, *p*=0.0007), but not 5-FU MIC (r=0.014, *p*=0.9233), further emphasizing the compound-specific antibacterial effects of these fluoropyrimidines. Consistent with these data, *E. coli* was able to convert CAP to 5-FU during *in vitro* growth (**Extended Data Fig. 1e**). More work is needed to elucidate the mechanisms responsible for gut bacterial CAP activation and the metabolic fate of this prodrug during *in vitro* and *in vivo* growth, building upon the recent identification of a gut bacterial enzyme capable of CAP deglycosylation^4^.

Next, we sought to gain insight into the mechanism of action of fluoropyrimidines against human gut bacteria, building on foundational studies in *C. elegans^6,25,26^* and testing their generalizability to CAP and FUDR. In mammalian cells, the primary target of fluoropyrimidines is thymidylate synthase^30^, an essential enzyme for DNA, RNA, and protein biosynthesis^31^. This mechanism of action is nutrient-dependent; excess uracil decreases drug sensitivity^32^ (**Extended Data Fig. 2a**). As expected, the *E. coli* MIC for all three fluoropyrimidines was higher in rich media and increased by uracil in a dose-dependent manner (**Extended Data Figs. 2b-d** and **Supplementary Table 3**). Transcriptional profiling (RNA-seq) enabled a more comprehensive view of the metabolic pathways impacted by fluoropyrimidines. *E. coli* was grown to mid-exponential phase and then exposed to sub-MIC levels of CAP and 5-FU under both aerobic (n=2 independent experiments) and anaerobic conditions (**Supplementary Table 4**). Despite marked differences in transcriptional activity between growth conditions and experiments (**Extended Data Fig. 3a**), we were able to identify significant effects of both drugs relative to vehicle controls (**Extended Data Figs. 3b,c**). Pathway enrichment analysis revealed that pyrimidine metabolism was consistently impacted by 5-FU during aerobic growth; whereas flagellar assembly was consistently impacted by CAP irrespective of growth condition and by 5-FU under anaerobic conditions (**Extended Data Fig. 3d)**. Across both experiments, we identified 1,720 differentially expressed genes (DEGs; FDR<0.1, |log2 fold-change|>1, DESeq; **Extended Data Fig. 3e** and **Supplementary Table 5**). In our repeat experiment, 620/892 (69.5%) DEGs were unique to a single condition. We also identified 112 (CAP) and 12 (5-FU) genes consistently differentially expressed in both growth conditions. CAP exposure downregulated the flagellar biosynthetic pathway, including the master flagellar regulator *flhDC* (**Extended Data Fig. 3f**). 5-FU exposure upregulated *carA* expression, a key gene involved in the first committed step of the *de novo* pyrimidine biosynthetic pathway. Consistent with these results and the prior literature^26,33^, a targeted screen of 22 non-essential DEGs involved in pyrimidine metabolism revealed that the deletion of *upp* (uracil phosphoribosyltransferase) leads to a high-level of resistance to CAP, 5-FU, and FUDR (**Extended Data Figs. 2e-g** and **Supplementary Table 3**). Targeted and whole genome sequencing of 5-FU-resistant mutants revealed multiple unique single nucleotide polymorphisms, deletions, and frameshifts within the *upp* genes of *E. coli* and *B. fragilis* (**Supplementary Table 2**). In strains with wild-type *upp,* we identified mutations within other pyrimidine metabolism genes, including uridine phosphorylase (*E. coli*), uridylate kinase (*E. coli*), and thymidine kinase (*B. ovatus*) (**Supplementary Table 2**). Together, our results support pyrimidine metabolism as a key target for fluoropyrimidines, while also demonstrating broader impacts of these compounds on genes involved in flagellar assembly, consistent with another recent report^26^.

Drug resistant bacteria can often catalyze drug inactivation^34,35^, potentially interfering with the efficacy of cancer therapy due to decreased bioavailability and/or enhanced clearance^36^. To test if the identified fluoropyrimidine resistant gut bacteria were capable of drug inactivation, we screened the top 23 strains (5-FU MIC ≥62.5 μg/ml) using a 5-FU disk diffusion assay. This concentration is physiologically relevant as indicated by the peak plasma concentration of 5-FU achieved during intravenous chemotherapy^37^. We identified two active Proteobacterial strains: *E. coli* MG1655 and *Salmonella enterica* LT2 (**Fig. 1b**). LC-QTOF/MS confirmed the depletion of 5-FU with near quantitative conversion to DHFU (**Fig. 1c**). In mammalian cells, dihydropyrimidine dehydrogenase (DPYD) is responsible for the biotransformation of 5-FU to the inactive metabolite DHFU^38–40^. Patients with DPYD deficiency or single nucleotide polymorphisms within *DPYD* are at risk for elevated systemic exposure to 5-FU and adverse events^38–40^. In *E. coli*, DPYD is encoded by two neighboring genes: *preT* and *preA* found within the *preTA* operon^41^. Purified PreTA protein is sufficient to catalyze the reduction of the pyrimidines uracil and thymine, as well as 5-FU^41^; however, the genes necessary and sufficient for this activity in bacterial cells remained unclear.

We generated a clean deletion of the *preTA* operon in *E. coli* MG1655 (Δ*preTA*), which led to a complete loss-of-function in our bioassay that was validated by LC-QTOF/MS (**Fig. 1d**). Chromosomal complementation with a constitutively overexpressed *preTA* operon (*preTA^++^*) resulted in an increased rate of metabolism relative to wild-type *E. coli* (**Fig. 1d**). An integrated empty vector control (*E. coli* Δ*preTA*/pINT1) had no impact on either assay. We confirmed the role of the *preTA* operon in 5-FU inactivation with a second strain of *E. coli* (BW25113; **Extended Data Fig. 4a**) and demonstrated that both genes of the operon were required for conversion to DHFU (**Extended Data Fig. 4b**). Comparable growth was observed for each isogenic strain in the presence or absence of 5-FU, with the exception of *preTA^++^*, which showed an increase in drug tolerance in minimal media (**Extended Data Fig. 4c**). *E. coli* MG1655 growth was unaffected by DHFU (MIC>1 mg/ml). To test the potential relevance of this reaction to drug efficacy, we added cell-free supernatants (CFS) from each strain to the colorectal cancer (CRC) cell line HCT-116. Sterile media and *preTA*-deficient CFS supplemented with 5-FU inhibited cell proliferation; however, this effect was fully rescued by wild-type and *preTA^++^ E. coli* (**Fig. 1e**). Taken together, these results demonstrate that *preTA* is both necessary and sufficient for 5-FU inactivation by *E. coli* (**Fig. 1f**), and that *preTA* is just one of many mechanisms that contribute to variations in drug sensitivity between gut bacterial strains.

We were surprised that our cell-based assays had demonstrated near quantitative conversion of 5-FU to DHFU given that *E. coli* PreTA has been characterized as a reversible enzyme^41^. To explore this finding in more detail, we turned to *in vitro* studies with *E. coli* PreTA (**Fig. 2a**). Purifying heterologously expressed *E. coli* PreTA to homogeneity showed that PreTA purifies as a heterotetramer and contains both flavin and iron-sulfur cofactors as seen by characteristic shifts in the oxidized and reduced UV– visible absorption spectra (**Fig. 2b, Extended Data Figs. 5a-c**). We next confirmed the activity of the heterologous PreTA by anaerobically incubating the protein with substrates and reducing equivalents, observing the concomitant decreases of substrate peaks with increases of product peaks by high pressure liquid chromatography (HPLC) (**Fig. 2c, Extended Data Fig. 5d**). The correct product masses of these reactions were identified by Liquid Chromatography High Resolution Mass Spectrometry (LC-HRMS), indicating that the protein was catalyzing the expected reaction (**Extended Data Fig. 5e**). Finally, we measured steady-state kinetics in both the reductive and oxidative reaction directions and saw that while the catalytic efficiency for uracil and 5-FU was comparable in the reductive direction, the catalytic efficiency in the oxidative direction decreased by two orders of magnitude for DHU compared to the reductive direction (**Figs. 2d,e** and **Supplementary Table 6**). While literature has stated DPYD is a reversible enzyme^42–44^, quantification of this reversibility indicates that the reaction proceeds to near completion in the reductive direction^45,46^. Together, these data suggest that under physiological conditions DHFU is an extremely poor substrate for PreTA and the metabolism of 5-FU by bacteria at physiologically relevant concentrations is essentially irreversible (**Fig. 2f**), consistent with our cell-based assay data (**Figs. 1c,d**).

**Figure 2.**
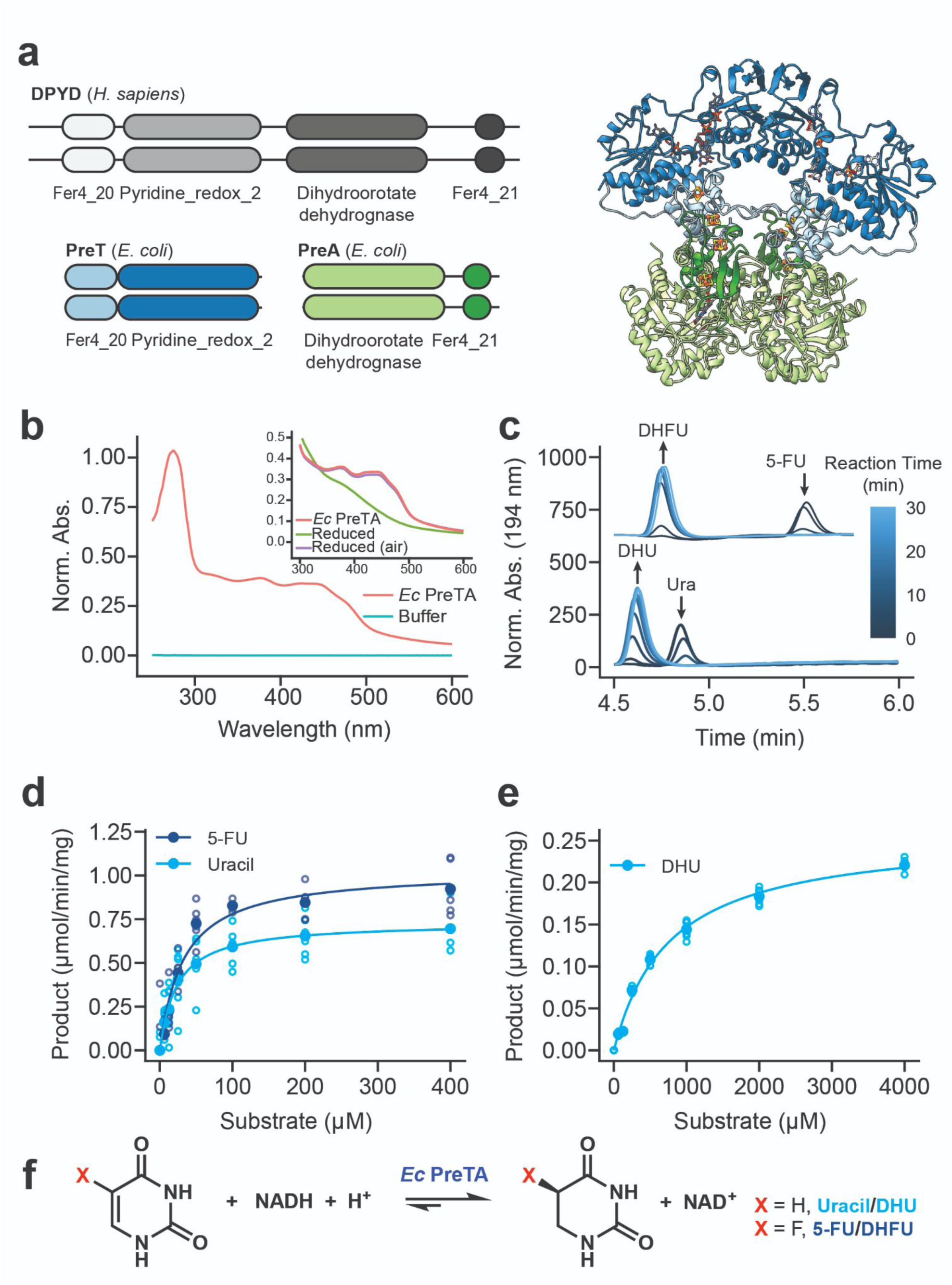
*E. coli* PreTA more rapidly catalyzes the reduction of pyrimidines. **(a)** A cartoon schematic shows the domain architecture of the homodimeric *H. sapiens* DPYD and heterotetrameric *E. coli* PreTA, as well as a *E. coli* PreTA homology model (Swiss-Model^57^). Protein backbone colors are indicated by the schematic, cofactors are colored by heteroatom. **(b)** UV–visible absorption spectra indicate heterologously expressed *E. coli* PreTA has characteristic peaks due to 4Fe-4S and flavin cofactors. Inset shows reduction of flavin cofactors and reoxidation upon exposure to air. **(c)** *In vitro* enzyme assays analyzed by high pressure liquid chromatography (HPLC) show conversion of uracil and 5-FU to their reduced forms over time. **(d-e)** Michaelis-Menten analysis of NADH cofactor consumption and accumulation *in vitro* enzyme assays in the reductive **(d)** and oxidative **(e)** directions display a marked difference in the ability for *E. coli* PreTA to catalyze the oxidative reaction. Solid points represent the mean of the hollow circles, n=6 replicates over 2 different enzyme preparations. **(f)** Schematic of the chemical reaction catalyzed by *E. coli* PreTA.

Measurements of tumor growth following CAP administration provided additional support for the physiological relevance of *E. coli* PreTA. We opted to use the *preTA^++^* and Δ*preTA E. coli* strains to avoid the potential confounding effects of transcriptional regulation; the wild-type *preTA* operon was expressed at a significantly higher level under aerobic conditions (**Extended Data Fig. 3g**). We used a tumor xenograft model to test the impact of PreTA on drug efficacy in three independent experiments that varied in the enrollment criteria and streptomycin dose (**Fig. 3a** and **Extended Data Figs. 6a,g**). Mice injected with HCT-116 human colorectal cancer cells were randomly split into two colonization (Δ*preTA* or *preTA^++^ E. coli*) and two treatment (CAP or vehicle) groups. As expected, baseline tumor volumes were highly variable in experiments utilizing a set enrollment date (**Extended Data Figs. 6b,h**), in contrast to rolling enrollment based on tumor size (**Fig. 3b**); there were no significant differences between groups. *E. coli* colonization level was dose-dependent and similar between groups (**Extended Data Figs. 6c,i** and **7a**). Mice colonized with high levels of Δ*preTA E. coli* that received CAP had a significant decrease in tumor growth relative to vehicle controls or CAP treated animals colonized with the *preTA^++^* strain (**Figs. 3c-e**, rolling enrollment) corresponding to significantly prolonged survival (**Fig. 3f**). Similar trends were observed in the high streptomycin, set enrollment experiment (**Extended Data Figs. 6d-f**); however, low streptomycin did not result in detectable differences between groups (**Extended Data Figs. 6j-l**). These results are consistent with the hypothesis that variations in the abundance of *preTA* encoding bacteria can lead to significant differences in drug efficacy.

**Figure 3.**
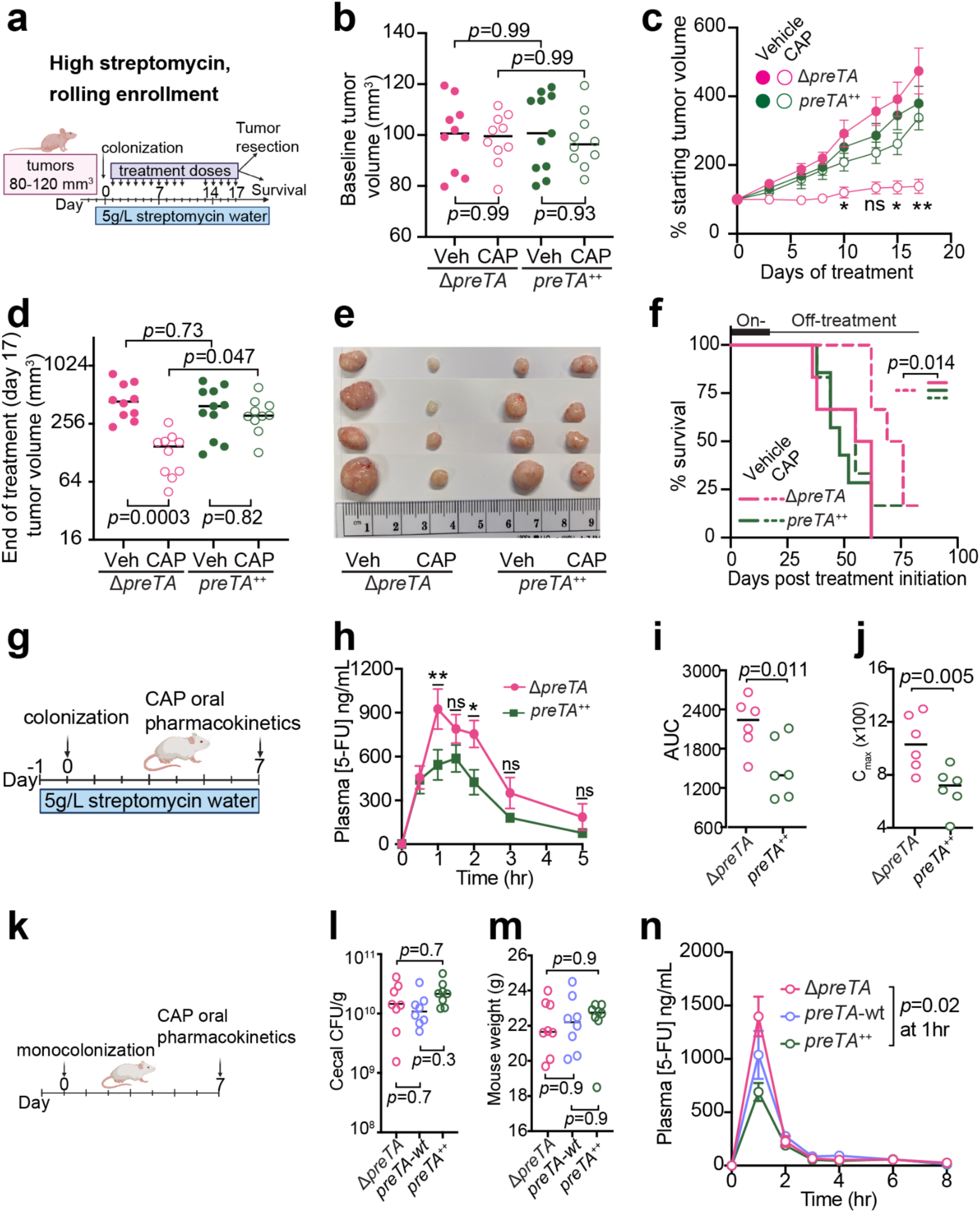
PreTA interferes with capecitabine (CAP) efficacy in mice. **(a)** HCT-116 human colorectal cancer cells were injected into 5-6-week-old female athymic nude mice (n=10-11 mice/group). When the tumor volume reaches 80-120 mm^3^, the mice were given streptomycin water and colonized with Δ*preTA* or *preTA^++^ E. coli* followed by oral administration of CAP (100 mg/kg) or vehicle (Veh). At the end of treatment on day 17, randomly selected mice (n=4/group) were sacrificed for tumor resection and the remaining mice (n=6-7/group) were allowed to reach the humane endpoint. **(b)** Baseline tumor volume prior to treatment (n=10-11 mice/group; lines represent the median). **(c)** Percentage of starting tumor volumes across time (n=10-11 mice/group; Tukey’s multiple comparison-corrected 2-way ANOVA tests with the *p*-values highlighted for pairwise comparison between Δ*preTA-*CAP *preTA^++^*-CAP groups in the graph; error bars represent standard error). 2-way ANOVA multivariate model shows significant interactions between experimental groups and time (groups *p*=0.0004; time *p*<0.0001; groups x time *p*<0.0001). **(d)** Tumor volumes at the end of treatment on day 17 post treatment initiation (n=10-11 mice/group; lines represent the median). 2-way ANOVA multivariate model shows significant interactions between treatment and colonization (treatment *p*=0.0004; colonization *p*=0.23; treatment x colonization *p*=0.01). **(e)** Resected tumors at the end of treatment on day 17 (n=4 mice/group). **(f)** Percentage of mice reaching the humane endpoint defined as tumor length >20 mm (n=6-7 mice/group, log-rank Mantel-Cox test comparing Δ*preTA-*CAP vs. all other groups). Two mice were censored as they did not reach the endpoint when the experiment ended on day 83. **(g)** Experimental design for our pharmacokinetics (PK) experiments in CONV-R mice. 8-10-week-old male BALB/c mice were given streptomycin water one day prior to colonization with Δ*preTA* or *preTA^++^ E. coli*. 7 days post colonization, 1100 mg/kg CAP was orally administered and longitudinal plasma samples were collected from each mouse. **(h)** LC-MS/MS quantification of plasma 5-FU (n=6 mice/group; error bars represent standard error; ANOVA mixed effect analysis with Sidak’s multiple comparison correction). **(i)** Area under curve (AUC) for plasma 5-FU (n=6 mice/group; lines represent the median; Welch’s one-tail t-test). **(j)** Peak plasma concentrations of 5-FU (n=6 mice/group; lines represent the median; Welch’s one-tail t-test). **(k)** Experimental design for PK experiments in gnotobiotic mice. 8-10-week-old germ-free female BALB/c mice were mono-colonized with Δ*preTA, preTA^++^*, or wild-type *E. coli.* 7 days post colonization, 500 mg/kg CAP was orally administered and longitudinal plasma samples were collected from each mouse (n=8 mice/group). **(l)** Colony-forming-unit (CFU) per gram of cecal content harvested at the end of the pharmacokinetics experiment (n=8 mice/group; lines represent the median). **(m)** Mouse weight measured on day 7 (n=8 mice/group; lines represent the median). **(n)** LC-MS/MS quantification of plasma 5-FU (n=8 mice/group; error bars represent standard error; ANOVA mixed effects analysis with Tukey’s correction). *p*-values are two-way **(b,d)** or one-way **(l,m)** ANOVA with Tukey’s correction. \**p*-value<0.05, \*\**p*-value<0.01.

Given the design of our experiments and prior *in vitro* data, the most parsimonious explanation for our *in vivo* results is that PreTA contributes to the first-pass metabolism of CAP leading to an effective decrease in the dose available to the tumor. Three independent pharmacokinetics experiments were performed to test the effect of PreTA on 5-FU bioavailability (**Figs. 3g** and **3k**). Conventionally-raised (CONV-R) specific pathogen free mice colonized with *preTA^++^ E. coli* had decreased plasma concentrations of 5-FU relative to Δ*preTA* controls following the oral administration of a high dose of CAP (1100 mg/kg, **Fig. 3h**). Drug bioavailability indicated by area under curve **(Fig. 3i**) and peak plasma concentration (**Fig. 3j**) was significantly lower in *preTA^++^* colonized mice relative to Δ*preTA* controls. Similar trends were observed when we repeated the experiment with lower CAP dose (500 mg/kg, **Extended Data Fig. 6m**). To remove the potential confounding effect of the background microbiota, we conducted a similar study using gnotobiotic mice **(Fig. 3k)**. Gnotobiotic mice mono-colonized with Δ*preTA, wt,* or *preTA^++^ E. coli* had comparable colonization level (**Fig. 3l**) and weight (**Fig. 3m**). Plasma 5-FU concentration was significantly decreased in *preTA^++^* colonized mice relative to Δ*preTA* controls; the *wt* strain had an intermediate phenotype that did not reach statistical significance (**Fig. 3n**). DHFU was undetectable in plasma from all experiments. An alternative hypothesis is that changes in uracil metabolism have off-target effects on the rest of the gut microbiota, leading to more indirect effects on tumor growth. We performed 16S rRNA gene sequencing on longitudinal stool samples collected prior to and during treatment with streptomycin and CAP (**Supplementary Table 7**). Gut microbial diversity and community structure were significantly altered over time (**Extended Data Figs. 7b-f**), consistent with prior data in the streptomycin model^47^. We did not detect any significant differences in community structure, diversity, or *E. coli* abundance between colonization groups (**Extended Data Figs. 7b-f**). Only 3/69 amplicon sequence variants (ASVs) were differentially abundant between groups (**Extended Data Figs. 7g**). While these data support the first-pass metabolism hypothesis, additional studies are warranted to assess additional alternative hypotheses such as PreTA-dependent changes in intestinal drug absorption and/or hepatic metabolism.

Our findings in the model gut bacterium *E. coli* provide a valuable *proof-of-principle* and a tractable model for future mechanistic studies; however, the bacterial species that encode *preTA* in cancer patients remained unclear. Given the conserved sequence and function of mammalian DPYD and bacterial PreTA we were surprised that only two moderately drug resistant strains (MIC=62.5 µg/ml) were identified in our screen for 5-FU inactivating bacteria. A tBLASTn search querying *E. coli* PreTA against the draft genomes of the other 22 tested strains only revealed a single pair of genomic loci from *S. enterica* (84% and 94% full length amino acid identity to PreT and PreA, respectively). No significant hits were found in the 24 drug sensitive strains. To identify other putative 5-FU inactivating strains, we built profile Hidden Markov Models (HMMs) of *preT* and *preA* and used them to search for orthologs across 9,082 bacterial isolate genomes. To minimize false positives due to the fact that both PreT and PreA are part of large protein families with a range of functions and substrate specificities^48^, we required that the *preT* and *preA* orthologs were adjacent in the genome and on the same strand. This analysis revealed 1,704 putative *preTA* operons from 406 species, mainly in Proteobacteria (1,507 operons from 290 species) and Firmicutes (172 operons from 92 species) (**Extended Data Fig. 8a** and **Supplementary Table 8**). This phylum-level distribution was similar for species with an estimated gut prevalence over 5% (n=23) and for species with a gut prevalence of 1% or lower (n=109)^49^. As expected, *preTA* was conserved in close relatives of *E. coli*, including other *Escherichia*, *Salmonella*, and *Citrobacter*. However, we also found operons in more distantly related Proteobacteria (*e.g., Oxalobacter formigenes*) and diverse Firmicutes: *Anaerostipes hadrus*, *Eubacterium hallii*, and *Lactobacillus reuteri*. These results were consistent in an independent analysis of 60,664 <u>m</u>etagenome-<u>a</u>ssembled <u>g</u>enomes (MAGs)^50^, resulting in the identification of 74 additional *preTA* operons, all from human gut MAGs, 89% of which were Firmicutes or Proteobacteria (**Extended Data Fig. 8b** and **Supplementary Table 9**).

We validated 5 putative *preTA* orthologs found in bacterial isolate genomes through a combination of heterologous expression and cell-based assays. Heterologous expression in the *E. coli* Δ*preTA* strain enabled us to validate *preTA* orthologs detected in two additional Proteobacteria (*S. enterica* LT2, *O. formigenes* ATCC35274) and the Firmicute *L. reuteri* DSM20016 using both our disk diffusion and LC-QTOF/MS assays (**Extended Data Fig. 9a** and **Supplementary Tables 10,11**). Incubation of *S. enterica* LT2 and two additional *preTA-*positive strains that we did not evaluate using heterologous expression supported the ability of the Firmicutes *Anaerostipes caccae* DSM14662 and *Clostridium sporogenes* DSM795 to metabolize 5-FU (**Extended Data Fig. 9b**). Interestingly, the *preTA* from *A. caccae* depleted 5-FU without producing any detectable DHFU, potentially due to further downstream metabolism or an alternative pathway. Together, these results demonstrate that bacterial PreTA has maintained its activity against 5-FU despite considerable phylogenetic and primary sequence divergence (66-93% similarity).

Having identified putative *preTA* orthologs, we set out to quantify *preTA* abundance in metagenomes from a CRC patient population. We re-analyzed 575 metagenomic samples from 179 individuals who were sampled in an existing case-control study^51^, using a conservative pipeline to improve specificity for the *preTA* operon. Briefly, we looked for nucleotide matches to the *preTA* operons we identified, using regions around non-*preTA* homologs of *preT* and *preA* as decoys; we additionally filtered out regions with uneven coverage across the operon. Read counts were converted to reads per kilobase of genome equivalents (RPKG) as previously described^52^. We detected *preTA* in 99% of samples and in 100% of individuals (**Fig. 4a**). *preTA* abundance varied over approximately two orders of magnitude, indicating substantial variation between individuals. In this study, the average *preTA* abundance in the CRC cohort was 62% that of controls (*p*=0.0075), though within-cohort variation remained much larger than differences between cases and controls (**Fig. 4b**). This suggests that *preTA* is near universally present in the human gut microbiome, yet also highly variable in abundance. To determine what the major bacterial taxa contributing to *preTA* abundance were in human subjects, we divided the total reads per sample proportionally among clades from the Genome Taxonomy Database^53^ according to best sequence matches. At the class level, Clostridia were the dominant source of *preTA* (**Fig. 4c**). Most of these reads typically came from an operational taxonomic unit (OTU) in the genus *Anaerostipes*. These results, together with the significant depletion for *Anaerostipes* in CRC cases^54^, likely explain the overall differences between cases and controls in *preTA* abundance (**Fig. 4b**). In a minority of samples, *preTA* reads were primarily from *E. coli*, *L. reuteri*, *R. bromii*, or an OTU called CAG-307 (a member of the *Acholeplasmataceae*). Thus, in most samples, the *preTA*s that were most abundant in metagenomes were closely related to the *preTA*s whose activity we verified experimentally (**Extended Data Figs. 9a,b**).

**Figure 4.**
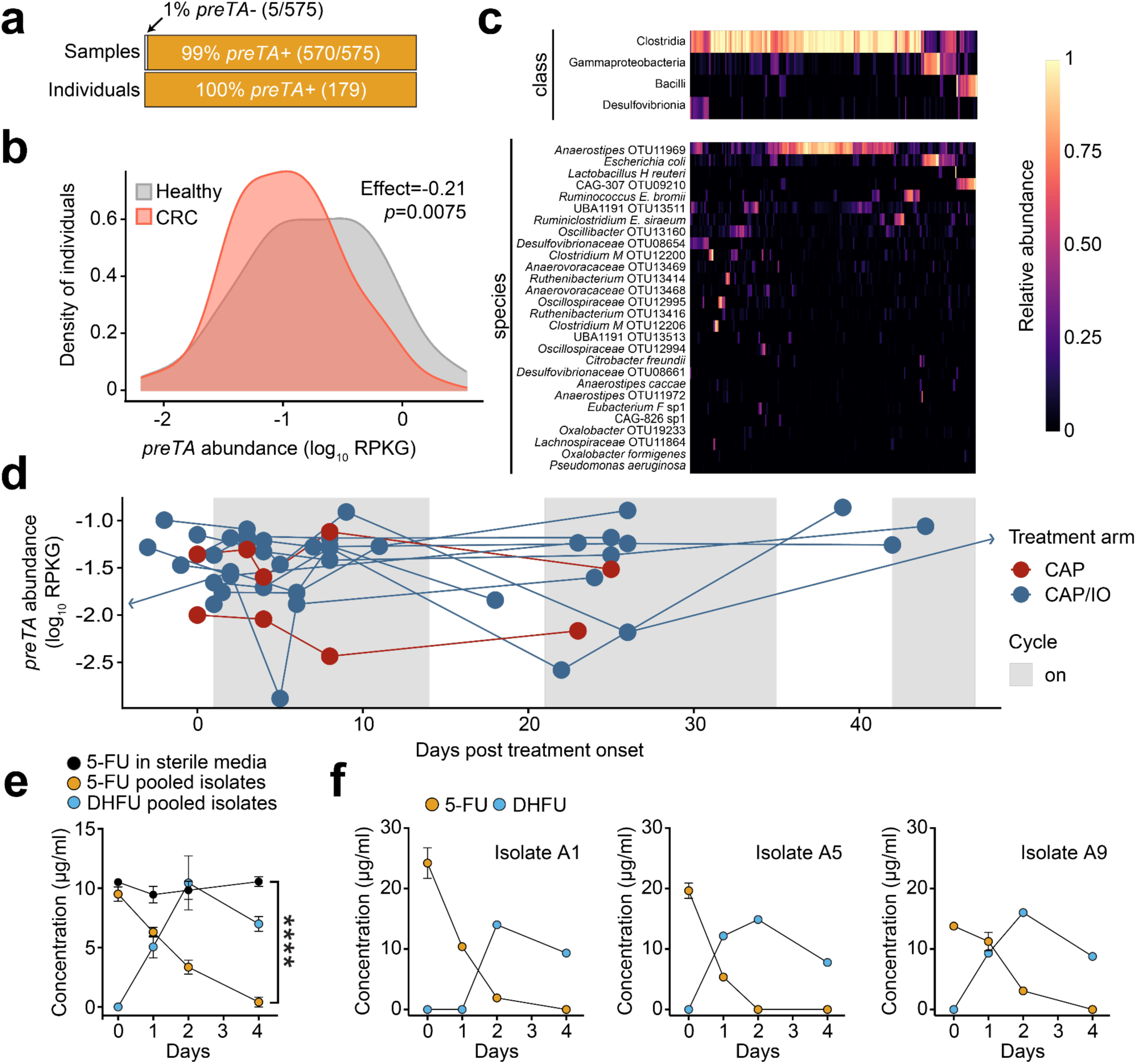
The *preTA* operon is prevalent and functional in the gut microbiomes of colorectal cancer (CRC) patients and controls, exhibiting marked inter-individual and temporal variation in abundance. **(a)** Number of samples and individuals where *preTA* was detected (orange) or not detected (light gray). **(b)** Variation in *preTA* abundance, as log_10_ reads per kilobase of genome equivalents (RPKG), across CRC patients and healthy controls where *preTA* was detected (n=179 individuals^51^). **(c)** Fraction of total reads mapping to *preTA* in CRC and control samples where it was detected (x-axis) whose best hit was in a given phylogenetic class (top) or species (bottom). Taxa are ordered by decreasing mean across all samples; samples are ordered based on hierarchical clustering using Bray-Curtis dissimilarity and complete linkage. **(d)** Abundance, as log_10_ RPKG, of the *preTA* operon in the gut microbiome prior to and during treatment with the oral fluoropyrimidine CAP (red) or a combination of CAP and immunotherapy (blue). Lines connect measurements for the same patient. One zero RPKG value was replaced with half the minimum non-zero value prior to taking the logarithm. 3/11 patients varied over an order of magnitude during treatment (**Extended Data Fig. 10**). The first day of treatment is defined as day 1. Two samples collected on days −16 and 66 were censored for display purposes. **(e)** Bacterial isolates from a representative CRC patient stool sample in the GO Study (patient 1, **Supplementary Table 12**) were isolated on McConkey agar, pooled, and incubated in BHI^+^ with 5-FU for 4 days and 5-FU/DHFU were quantified by LC-QTOF/MS. Values are mean±sem. \*\*\*\**p*-value<0.0001, two-way ANOVA contrasting 5-FU levels in sterile and inoculated media. **(f)** Three bacterial isolates from the same CRC patient stool sample were incubated as described in panel e (n=3 biological replicates). Values are mean±sem.

Finally, we examined the dynamics of *preTA* abundance over time in CRC patients treated with fluoropyrimidines. Stool samples were collected as part of the <u>G</u>ut Microbiome and <u>O</u>ral Fluoropyrimidine Study in Patients with Colorectal Cancer (GO; ClinicalTrials.gov NCT04054908). At the time of our pilot sequencing efforts, longitudinal samples collected before, during, and after the first treatment cycle were available from 11 patients treated with CAP-based regimens. In total, 54 samples were subjected to deep metagenomic sequencing (**Supplementary Table 12**). Consistent with the larger study of CRC patients and healthy controls (**Fig. 4b**), we detected marked variation in *preTA* abundance between individuals (**Fig. 4d**) and *preTA* reads in most individuals mapped best to *Anaerostipes* (**Extended Data Fig. 10**). In these initial 11 patients, we did not observe any consistent and statistically significant shifts in *preTA* abundance over time (*p*=0.42, linear model with fixed time effects and random patient effects, Satterthwaite approximation; **Fig. 4d**). In 3 individuals, *preTA* abundance varied over an order of magnitude across time points. Incubations of pooled bacterial isolates from the baseline samples of an expanded set of 22 CRC patients revealed 5-FU inactivation in 17/22 (77%) individuals (**Extended Data Fig. 9c**). LC-QTOF/MS analysis of incubations of pooled (**Fig. 4e**) and clonal (**Fig. 4f**) bacterial isolates from a representative patient confirmed the metabolism of 5-FU to DHFU. Taken together, these results suggest that variations in baseline *preTA* abundance and/or temporal fluctuations may contribute to differences in treatment outcomes.

Our results demonstrate how pathways for sensitivity to and metabolism of fluoropyrimidine drugs are conserved across two domains of life. These findings have broad implications across multiple traditional disciplines. Elucidating drug mechanisms of action in bacteria may provide translational insights for host tissues, raising questions as to the broader view of the off-target effects of therapeutics and the degree to which drug-induced shifts in microbial community structure and function have downstream consequences for drug efficacy and/or toxicity. While our *in vitro* and *in vivo* results both support the hypothesis that bacterial drug inactivation in the gastrointestinal tract or even within diseased tissue^5^ could interfere with drug efficacy, more work is needed to assess the relative impact of this biotransformation on efficacy versus gastrointestinal or other side effects, as previously demonstrated for the anti-cancer drug irinotecan^8^. Continued biochemical and structural characterization of the PreTA holoenzyme could enable the development of bacteria-specific enzyme inhibitors, while insights into the ecological role of PreTA may unlock strategies for strain replacement prior to therapy. These results are also consistent with another recent study^9^, which highlighted the unexpected overlap between host and bacterial drug metabolites. Traditional approaches for studying drug disposition do not distinguish these two alternatives^36^, which may explain the difficulties in predicting fluoropyrimidine toxicity using only human genotypic information. Furthermore, our discovery of diverse *preTA* positive bacterial strains could open the door towards preventing the severe, and at times lethal, toxicity observed from patients undergoing fluoropyrimidine chemotherapy with loss-of-function mutations in mammalian DPYD^38^.

Correspondence and requests for materials should be addressed to Peter J. Turnbaugh.

## Acknowledgements

We are indebted to the other members of the Gerona, Goga, Pollard, and Turnbaugh labs for their helpful suggestions during the preparation of this manuscript. We thank Brian Yu, Michelle Tan, and Rene Sit from the Chan-Zuckerberg Biohub for assistance with DNA sequencing; Gerry Wright and Linda Ejim for providing pINT1; Jordan Bisanz for advice with computational methods; Kyle Spitler in the UCSF Quantitative Metabolite Analysis Center for help with analytical methods; and the GO Study clinical research coordinators (Dalila Stanfield, Paige Steiding, and Julia Whitman). This work was supported by the National Institutes of Health (R01HL122593; R21CA227232; R01CA223817) and the Searle Scholars Program (SSP-2016-1352) and CDMRP W81XWH-18-1-0713. P.J.T. is a Chan Zuckerberg Biohub Investigator and was a Nadia’s Gift Foundation Innovator supported, in part, by the Damon Runyon Cancer Research Foundation (DRR-42-16). A.G. was in part supported by a MARK Foundation Endeavor Award. Fellowship support was provided by the Canadian Institutes of Health Research (P.S. and K.N.L.) and the NIH (J.V.L. - F32CA243548, T32CA108462; T.S.K. - F30CA257378). B.G.H.G. is a Connie and Bob Lurie Fellow of the Damon Runyon Cancer Research Foundation (DRG-2450-21).

## Author contributions

P.J.T conceived of the project and was the primary supervisor for the study. R.R.G., A.G., and K.S.P. also supervised components of this work. T.S.K. led the pharmacokinetics, xenograft, transcriptomics, and amplicon sequencing data generation and analysis. P.S. led the *in vitro* screens and *E. coli* strain construction. B.G.H.G. led the biochemical characterization of PreTA. P.H.B. led the bioinformatic analysis of *preTA* operons across genomes and microbiomes. J.M., Y.N.A.M., T.S.K., B.G.H.G., and M.S. performed mass spectrometry. K.N.L. sequenced and analyzed the drug resistant *E. coli* mutants. J.V.L. assisted with the tumor xenograft measurements. C.E.A., A.V., and W.K. (GO Study PI) oversaw the conception and design of the GO Study and contributed patient samples. E.L.V.B. contributed to developing the study protocol and supervision of data collection for the GO Study. D.G. designed the GO Study Specimen Collection Kits and managed biospecimen collection, storage, and retrieval. P.S. wrote the initial draft. T.S.K. and P.J.T. revised the manuscript with input from all authors.

## EXTENDED DATA FIGURES AND LEGENDS

**Extended Data Figure 1.**
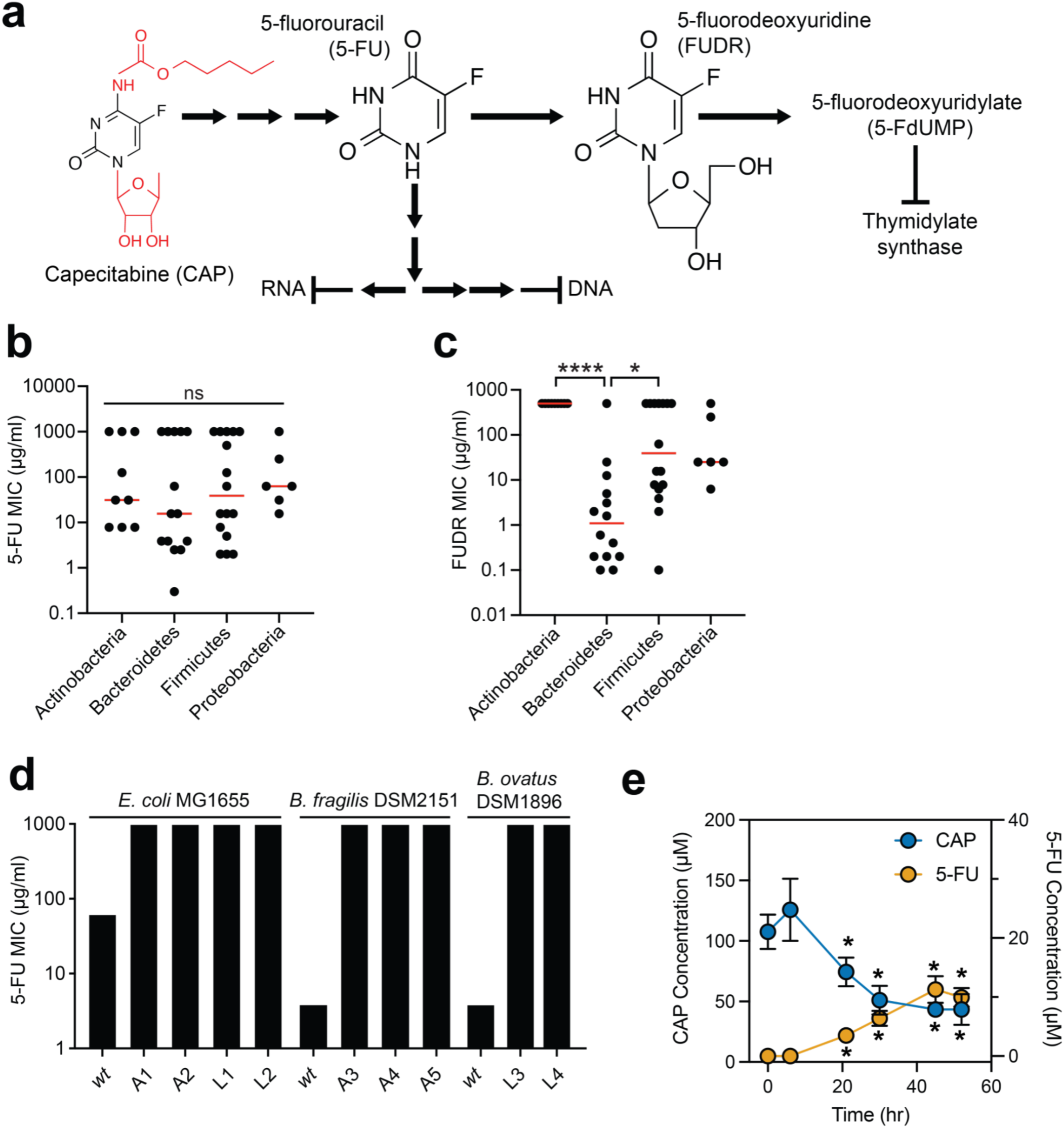
Bacterial taxa vary in their sensitivity to fluoropyrimidines despite the rapid emergence of resistance during *in vitro* growth. **(a)** Simplified metabolic pathway for the bioactivation of the oral prodrug capecitabine (CAP) to 5-fluorouracil (5-FU), 5-fluorodeoxyuridine (FUDR), and downstream metabolites^58^. Red indicates chemical groups hydrolyzed during the conversion of CAP to 5-FU. Sequential reactions are indicated by multiple arrows. **(b,c)** Variation in **(b)** 5-FU and **(c)** FUDR MIC at the phylum level. Bacterial strains where no MIC could be determined were set to the maximum tested concentration. Each dot represents a bacterial isolate. Red lines indicate the median. \**p*-value<0.05, \*\*\*\**p*-value<0.0001, Kruskal-Wallis test with Dunn’s correction for multiple comparisons. **(d)** 5-FU MICs of parent and 5-FU-resistant strains of *E. coli* MG1655, *B. fragilis* DSM2151, and *B. ovatus* DSM1896 (see **Supplementary Table 2**). The type of selection is indicated by the strain identifier: A=agar; L=liquid. **(e)** Wild-type *E. coli* BW25113 was assayed for conversion of CAP to 5-FU by LC-MS/MS (n=3 biological replicates per group). \**p*-value<0.05, 2-way ANOVA with Tukey’s test for multiple correction compared to the 0 hr time point of the same analyte.

**Extended Data Figure 2.**
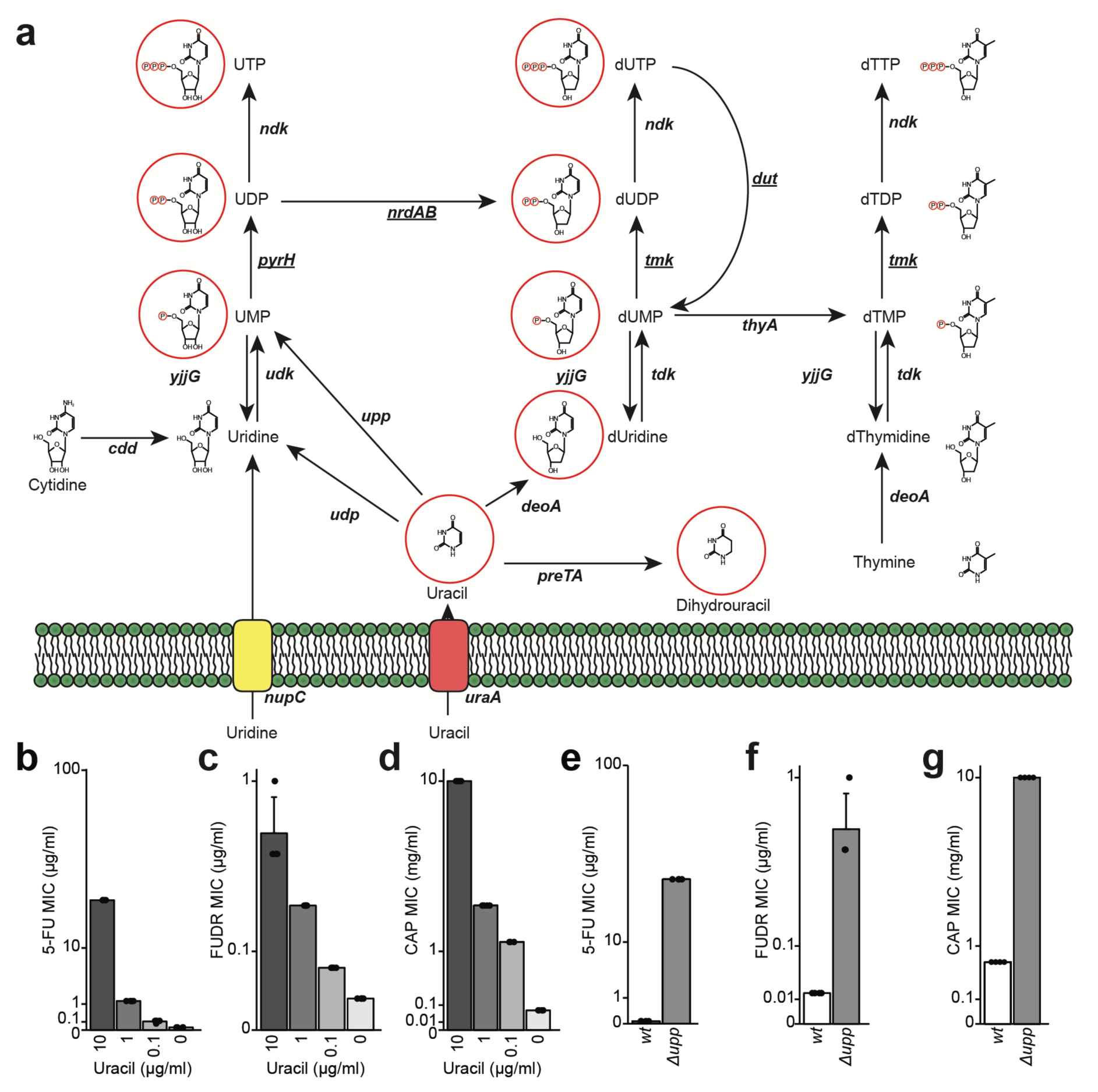
Multiple fluoropyrimidines disrupt the pyrimidine metabolism pathway in *E. coli*. **(a)** A working model for uracil and uridine import and metabolism in bacterial and mammalian cells. Underlined genes are essential for *E. coli* growth and metabolites with red circles indicate putative 5-FU metabolites. **(b-d)** Uracil rescues the growth of *E. coli* BW25113 in the presence of **(b)** 5-FU, **(c)** FUDR, and **(d)** CAP in a dose-dependent manner. M9MM plus glucose was used as the base media with added uracil from 0.1-10 µg/ml. **(e-g)** A loss-of-function mutation in uracil phosphoribosyltransferase gene (Δ*upp*) rescues the growth of *E. coli* BW25113 in the presence of **(e)** 5-FU, **(f)** FUDR, and **(g)** CAP. MIC assays performed in M9MM plus glucose. Values in panels b-g are mean±stdev (n=4 biological replicates per group).

**Extended Data Figure 3.**
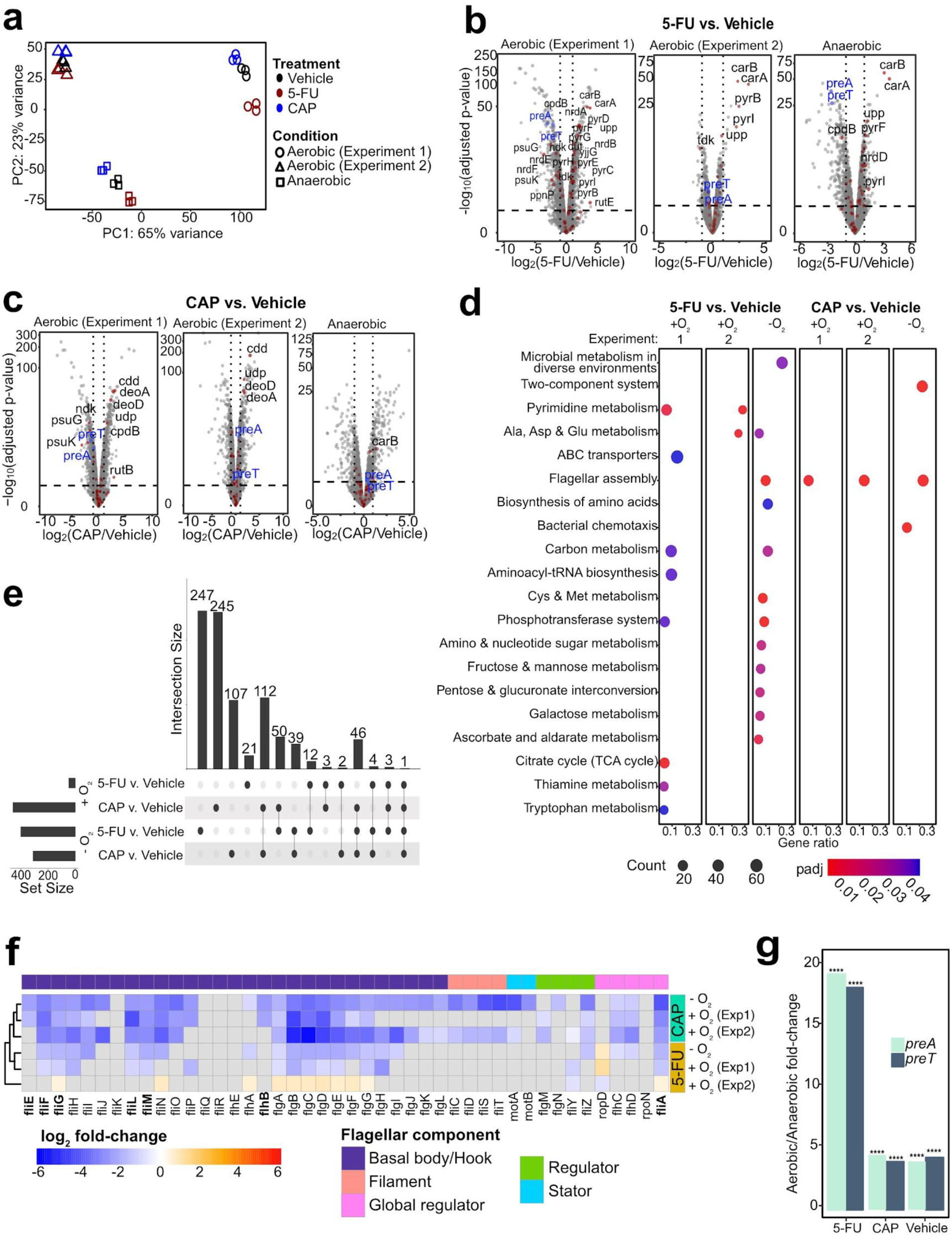
Shared and unique transcriptional response to the related fluoropyrimidines 5-fluorouracil (5-FU) and capecitabine (CAP) during aerobic and anaerobic growth. **(a)** Principal component analysis of *E. coli* MG1655 transcriptomes under different treatments in aerobic and anaerobic conditions (n=3 biological replicates; 2 independent experiments were conducted for aerobic growth). **(b,c)** Volcano plots of differentially expressed genes (DEGs; FDR<0.1, |log2 fold-change|>1). DEGs involved in pyrimidine metabolism are labeled by gene name and marked with a red dot. *preT* and *preA* are labeled in blue. **(d)** Pathway enrichment analysis following 5-FU and CAP exposure under aerobic and anaerobic conditions. The Kyoto Encyclopedia of Genes and Genomes (KEGG) database was used to test for pathway enrichment (*q*-value<0.1, Benjamini-Hochberg correction, enrichKEGG function in clusterProfiler^59^). **(e)** Upset plot comparing the fluoropyrimidine responsive DEGs under different growth conditions (**Supplementary Table 5**). **(f)** Heatmap of transcriptional changes of 44 genes involved in flagellar biosynthesis in response to fluoropyrimidine treatments under anaerobic and anaerobic conditions. Gray boxes indicate non-significant changes (FDR>0.1). Bolded genes indicate DEGs that are down-regulated in 5-FU evolved strains reported in literature ^26^. **(g)** *preT* and *preA* transcript levels are significantly lower during anaerobic growth relative to aerobic growth irrespective of the presence of fluoropyrimidines (****FDR<10^-27^, DESeq).

**Extended Data Figure 4.**
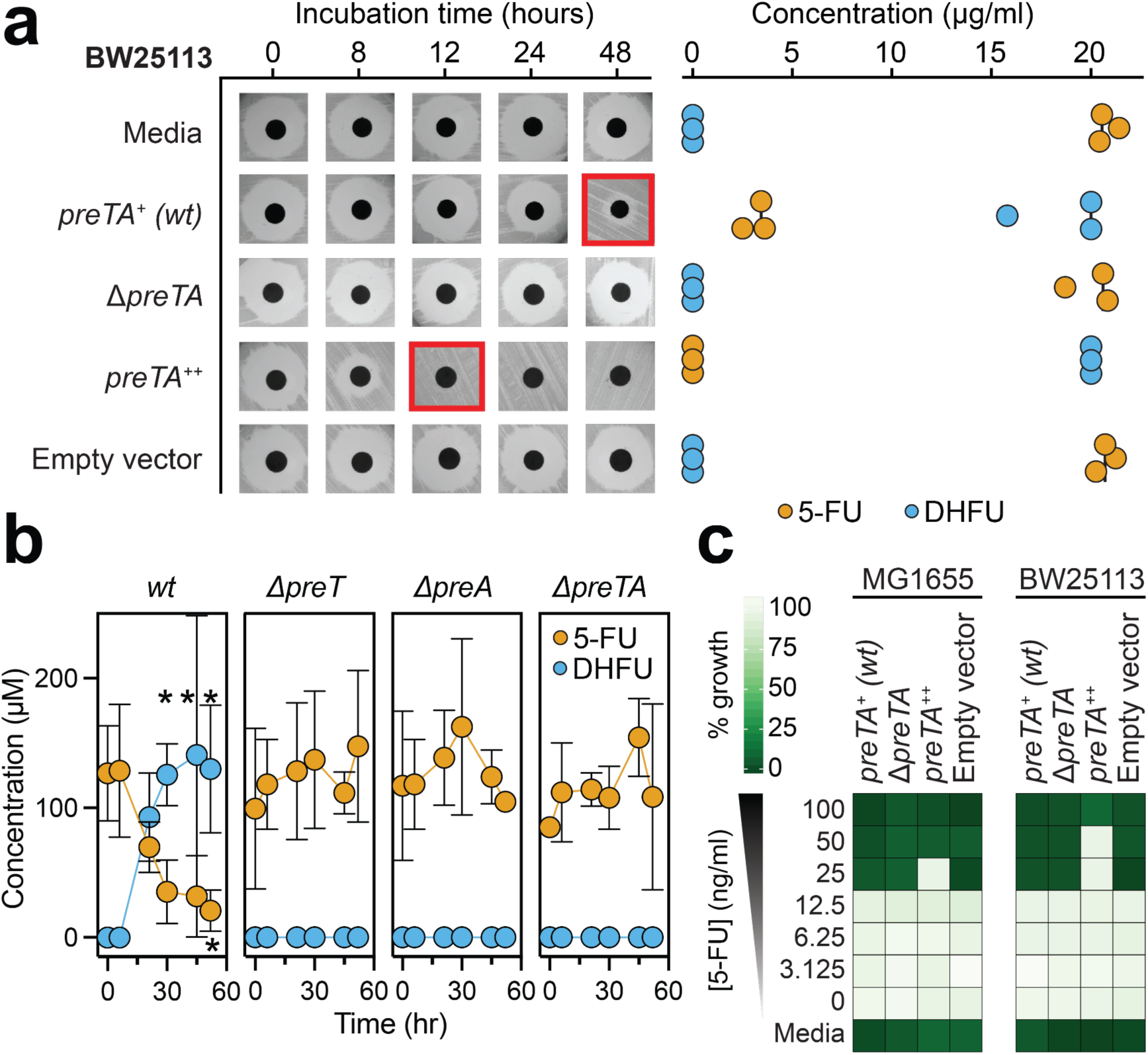
The *preTA* operon is necessary and sufficient for the inactivation of 5-fluorouracil in an independent *E. coli* background and has a modest impact on growth. **(a)** Wild-type (*preTA^+^*), deletion (*ΔpreTA*), complemented (*preTA^++^*), and empty vector *E. coli* BW25113 strains were assayed for residual 5-FU using disk diffusion (0-48 hours incubation) and LC-QTOF/MS (48 hours). Complementation and empty vector were on the *ΔpreTA* background. **(b)** Wild-type, single gene deletion (*ΔpreT*, *ΔpreA*) and operon deletion (*ΔpreTA*) strains of *E. coli* BW25113 were assayed for conversion of 5-FU to DHFU by LC-MS/MS (n=3, error bars represent standard deviation). \**p*-value<0.05, 2-way ANOVA with Tukey’s test for multiple correction compared to the 0 hr time point of the same analyte. **(c)** 5-FU MIC determination of the *E. coli* strains shown in panel a and Fig. 1d in minimal (M9MM) media. Values are normalized to the growth control (no 5-FU) with darker colors indicating growth inhibition. Sterile media is shown in the final row.

**Extended Data Figure 5.**
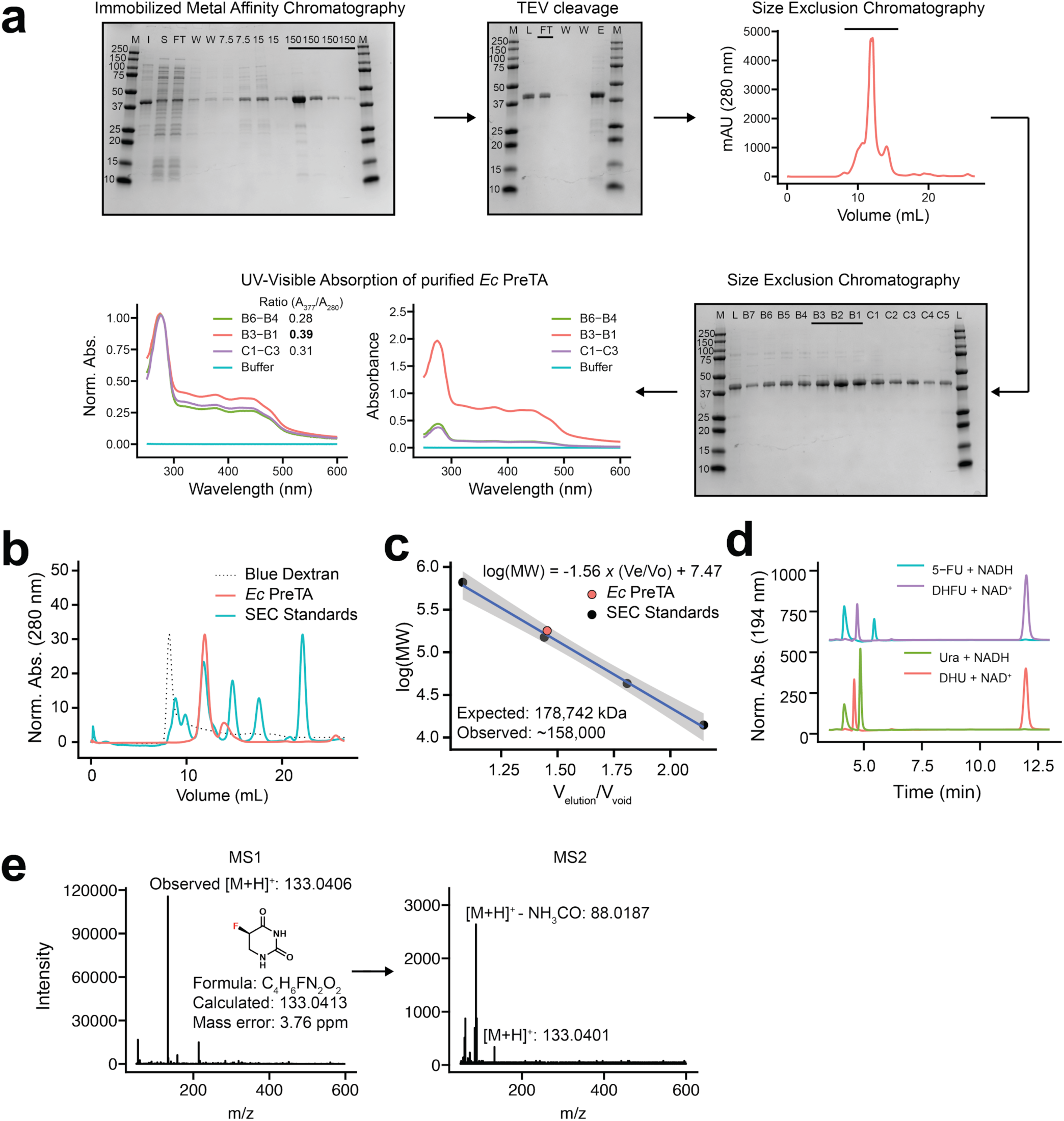
Purification and biochemical characterization of *E. coli* PreTA. **(a)** Purification workflow for *E. coli* PreTA including immobilized metal affinity chromatography (M: marker, I: insoluble fraction, S: soluble fraction, FT: flowthrough, W: wash, numbers across the top indicate imidazole concentration), TEV cleavage (M: marker, L: loaded, FT: flowthrough, W: wash, E: elute), size exclusion chromatography (M: marker, L: loaded, lanes are labeled by fraction from a 96 well plate collected via FPLC), and by UV–visible absorption spectroscopy where normalizing protein levels showed a ratio of greater than 0.35 for A_280_/A_377_ indicated holoenzyme. In all gels, numbers down the side indicate protein molecular weight in kDa and solid lines indicate lanes that were carried forward in the preparation. **(b-c)** Analytical size exclusion chromatography (SEC) traces **(b)** and analysis **(c),** characterizing the main peak as a heterotetramer. **(d)** High pressure liquid chromatography (HPLC) of NADH, uracil (Ura), dihydrouracil (DHU), NAD^+^, 5-fluorouracil (5-FU), and dihydrofluorouracil (DHFU). **(e)** Liquid Chromatography Mass Spectrometry (LC-MS/MS) of enzymatic reaction with 5-FU confirms the presence of the exact mass of DHFU.

**Extended Data Figure 6.**
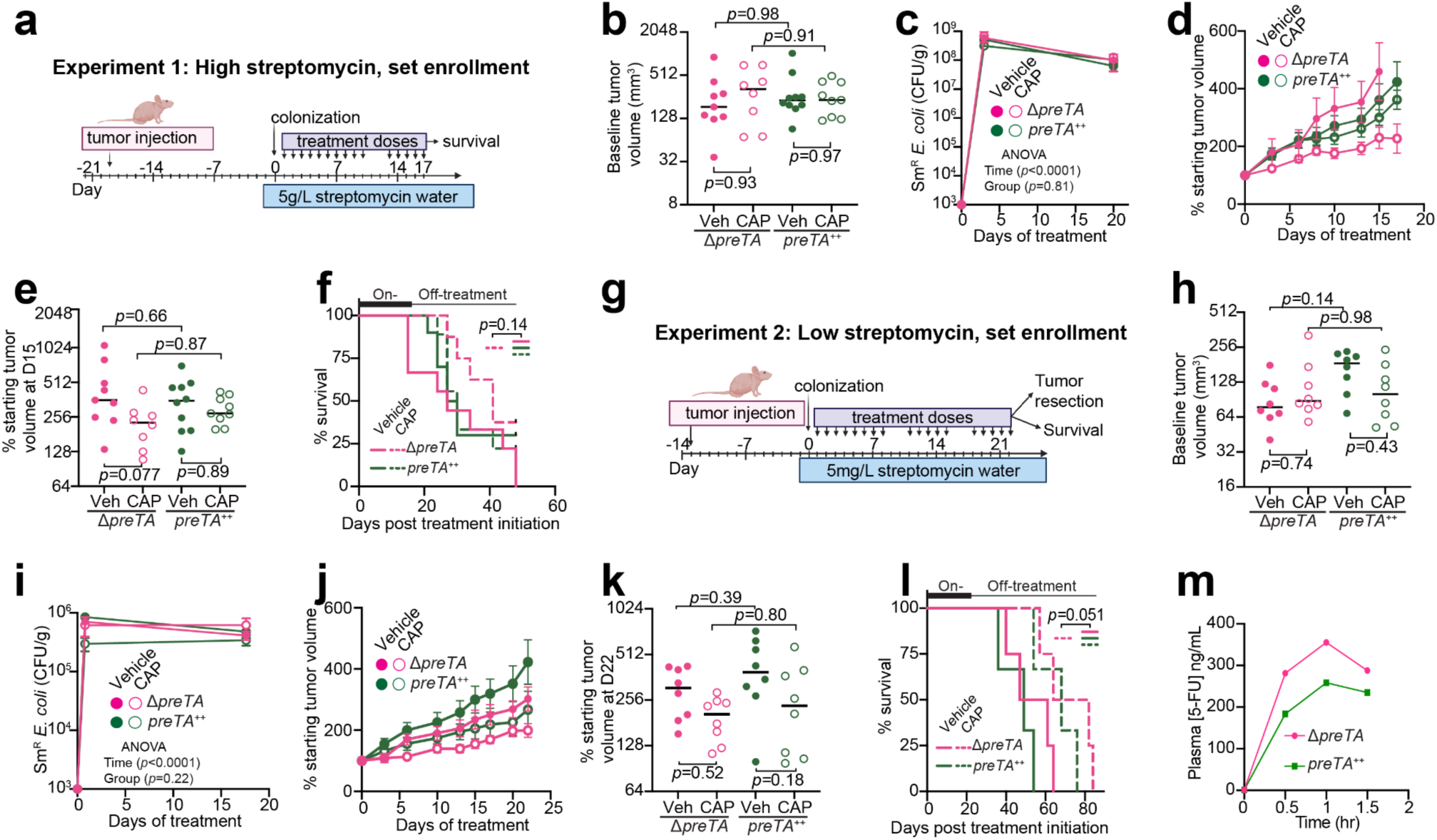
PreTA decreases the efficacy and oral bioavailability of capecitabine (CAP). **(a, g)** Experimental designs for our xenograft experiments 1 and 2, respectively. **(b, h)** Baseline tumor volumes prior to treatment (n=8-10 mice/group for experiment 1, n=8 mice/group for experiment 2; lines represent the median). **(c, i)** Quantification of streptomycin-resistant *E. coli* in feces across time (n=6 mice/group for experiment 1, n=8 mice/group for experiment 2, 2-way ANOVA with Tukey’s multiple comparison correction). No colonies were detected at time zero. During the log transformation, we replaced the zeros with 1000 CFU/g, which is our limit of detection. **(d, j)** Percentage of starting tumor volumes over time (n=8-10 mice/group for experiment 1, n=8 mice/group for experiment 2; error bars represent standard error). **(e, k)** Percentage of starting tumor volumes on **(e)** day 15 (n=8-10 mice/group) and **(k)** day 22 (n=8 mice/group); timepoint selected to capture all mice prior to euthanasia; lines represent the median. **(f, l)** Percentage of mice reaching the humane endpoint defined as tumor length >20 mm (n=8-10 mice/group for experiment 1, n=3-4 mice/group for experiment 2, Log-rank Mantel-Cox test comparing Δ*preTA-*CAP to all other groups). In experiment 1, 8 mice that did not reach the endpoint when the experiment ended at day 48 were censored. Censored mice were indicated by black boxes in the graph. *p*-values in panels (**b, e, h, k**) are from two-way ANOVA tests with Tukey’s multiple comparison corrections. **(m)** 8-10-week-old male BALB/c mice were given streptomycin water one day prior to colonization with Δ*preTA* or *preTA^++^ E. coli*. 7 days post colonization, 500 mg/kg CAP was orally administered and longitudinal plasma samples were collected from each mouse (see experimental design in Fig. 3g). LC-QTOF/MS was used to quantify 5-FU from pooled plasma samples following 500 mg/kg CAP administration (n=5 mice/group).

**Extended Data Figure 7.**
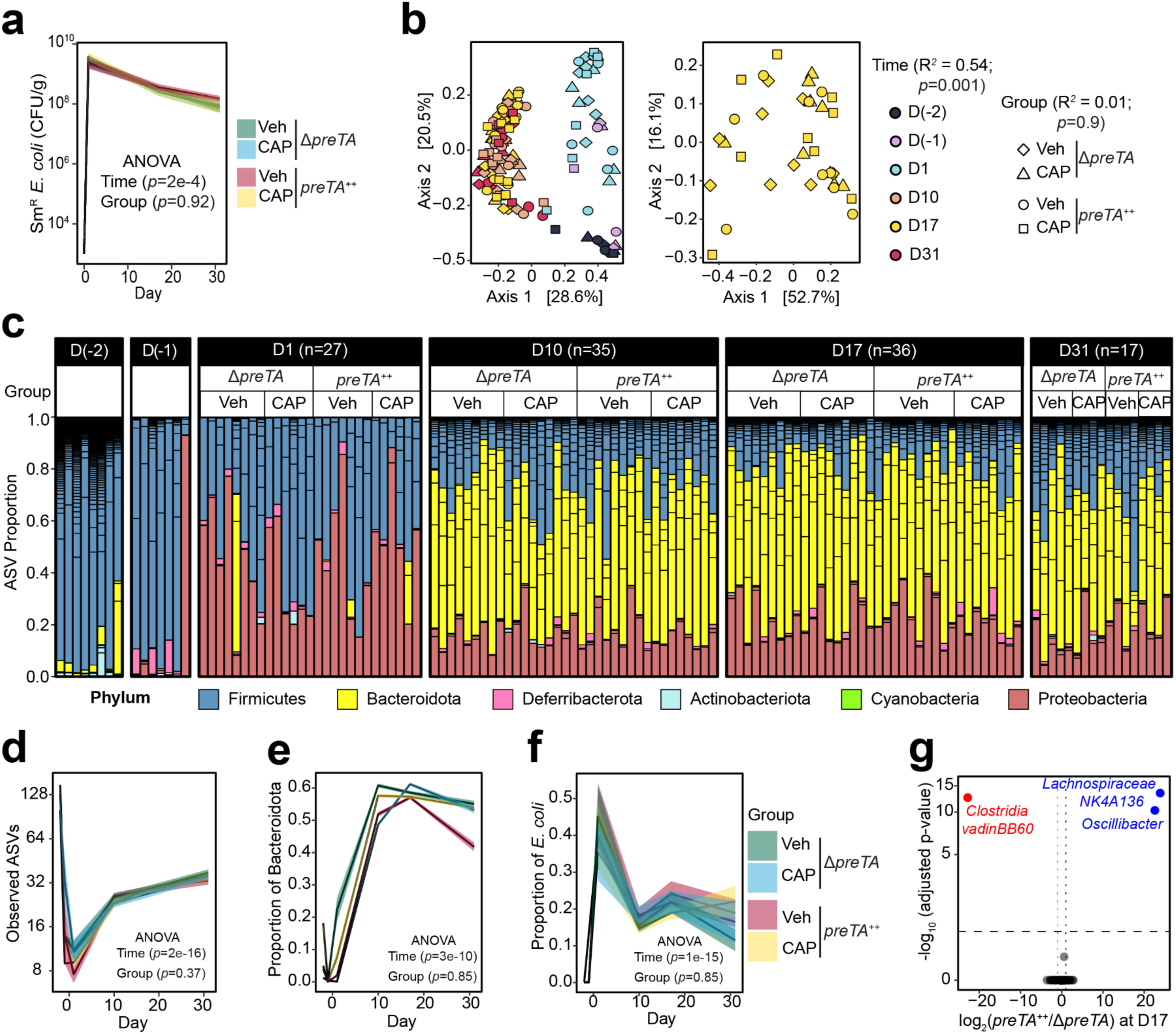
Consistent shifts in the gut microbiota across groups. **(a)** Quantification of streptomycin-resistant *E. coli* in feces across time (n=10-11 mice/group, 2-way ANOVA test). During log transformation, we replaced the zeros with 1000 CFU/g, which is our limit of detection. **(b)** Principal coordinate analysis of fecal microbiota from mice treated with CAP or vehicle (Veh) and colonized with Δ*preTA* or *preTA^++^ E. coli* across time (Bray-Curtis distance matrix, permutational multivariate analysis of variance test using ADONIS statistical package). **(c)** Microbial community composition at the phylum level. Each bar represents stool from each mouse. Short horizontal lines within bars represent different sequence variants. **(d)** Number of sequence variants observed over time for each treatment group (n=5-9 mice/group/time). **(e)** Proportion of Bacteroidota over time for each treatment group (n=5-9 mice/group/time). **(f)** Proportion of *E. coli* over time for each treatment group (n=5-9 mice/group/time). *p*-values in panels e-g are from 2-way ANOVA tests. **(g)** Volcano plots of differentially abundant sequence variants at day 17 (FDR<0.1, |log2 fold-change|>1).

**Extended Data Figure 8.**
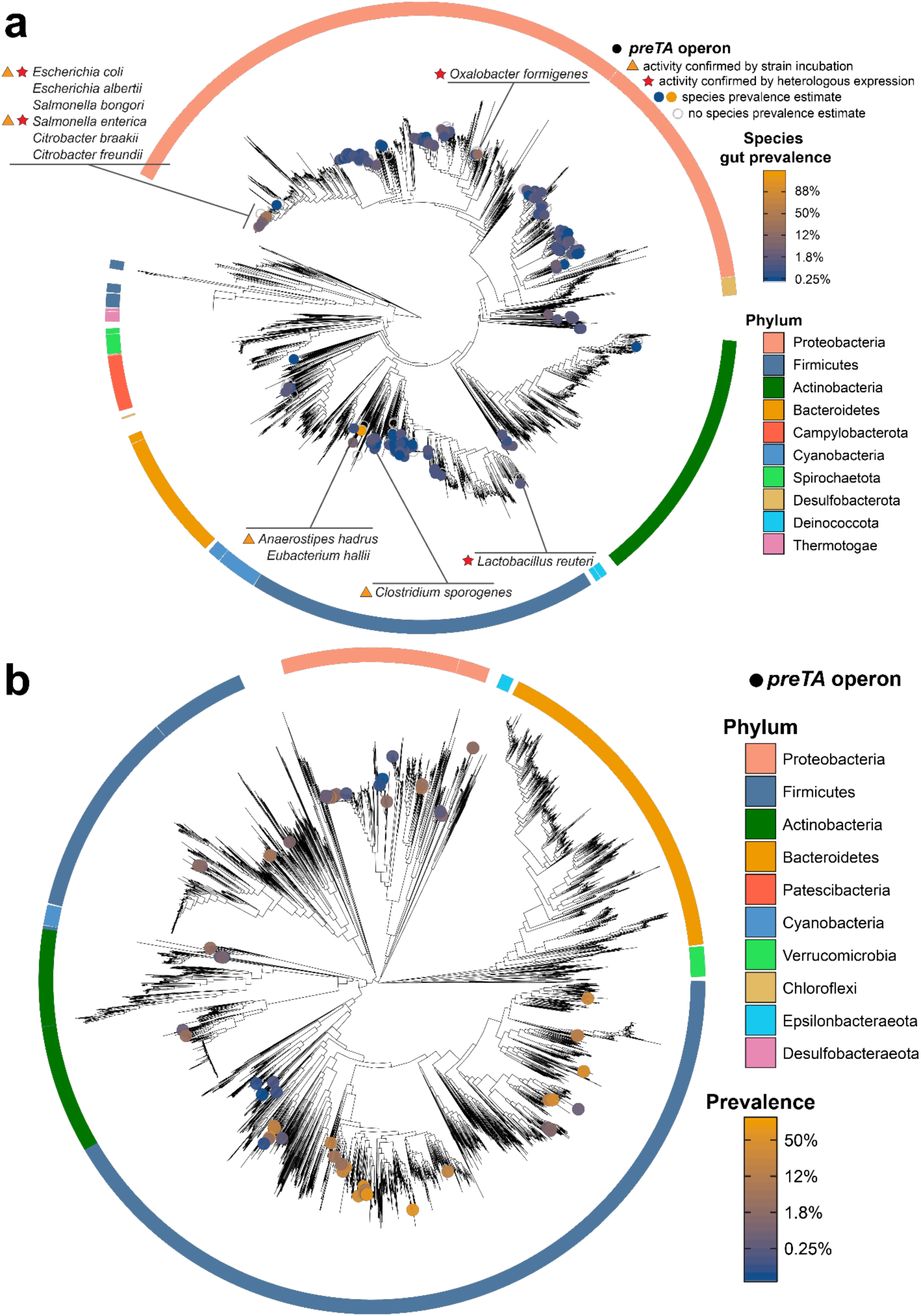
Functional orthologs of the *preTA* operon are widespread in human gut bacterial strains from the Firmicutes and Proteobacteria phyla. **(a)** Distribution of bioinformatically-identified *preTA* operons across RefSeq bacterial isolate genomes. A phylogenetic tree of these genomes made using a concatenated alignment of single-copy marker genes is shown. Bacterial species identified as carriers of putative *preTA* operons are identified with colored circles, where the color of the circle corresponds to prevalence levels from human gut microbiomes (blue: low prevalence; orange: high prevalence; unfilled grey: no prevalence estimate). Phylum-level annotations are shown as colored ring segments surrounding the tree for the ten phyla with the most species in Refseq. Specific taxa of interest are highlighted in call-out boxes. Red stars indicate *preTA* operons that have been validated to inactive 5-FU by heterologous expression from *E. coli* Δ*preTA* (see **Extended Data Fig. 9a**). Orange triangles indicate *preTA*-positive bacterial species (or close relatives) for which we have confirmed 5-FU inactivation *in vitro* (see **Extended Data Fig. 9b**). A list of *preTA*-positive bacteria can be found in **Supplementary Table 8. (b)** Distribution of putative *preTA* operons across gut metagenome-assembled genomes (MAGs). A phylogenetic tree of species from IGGdb, made using a concatenated alignment of single-copy marker genes, is shown in black. Species in which a MAG contains a putative *preTA* operon are identified with colored circles, where the color of the circle corresponds to prevalence in the gut (blue: low prevalence; orange: high prevalence). All species displayed were detected at least once from 3,810 gut metagenome samples. Phylum-level annotations are shown as colored ring segments surrounding the tree for the ten phyla with the most gut MAGs. Data can be found in **Supplementary Table 9**.

**Extended Data Figure 9.**
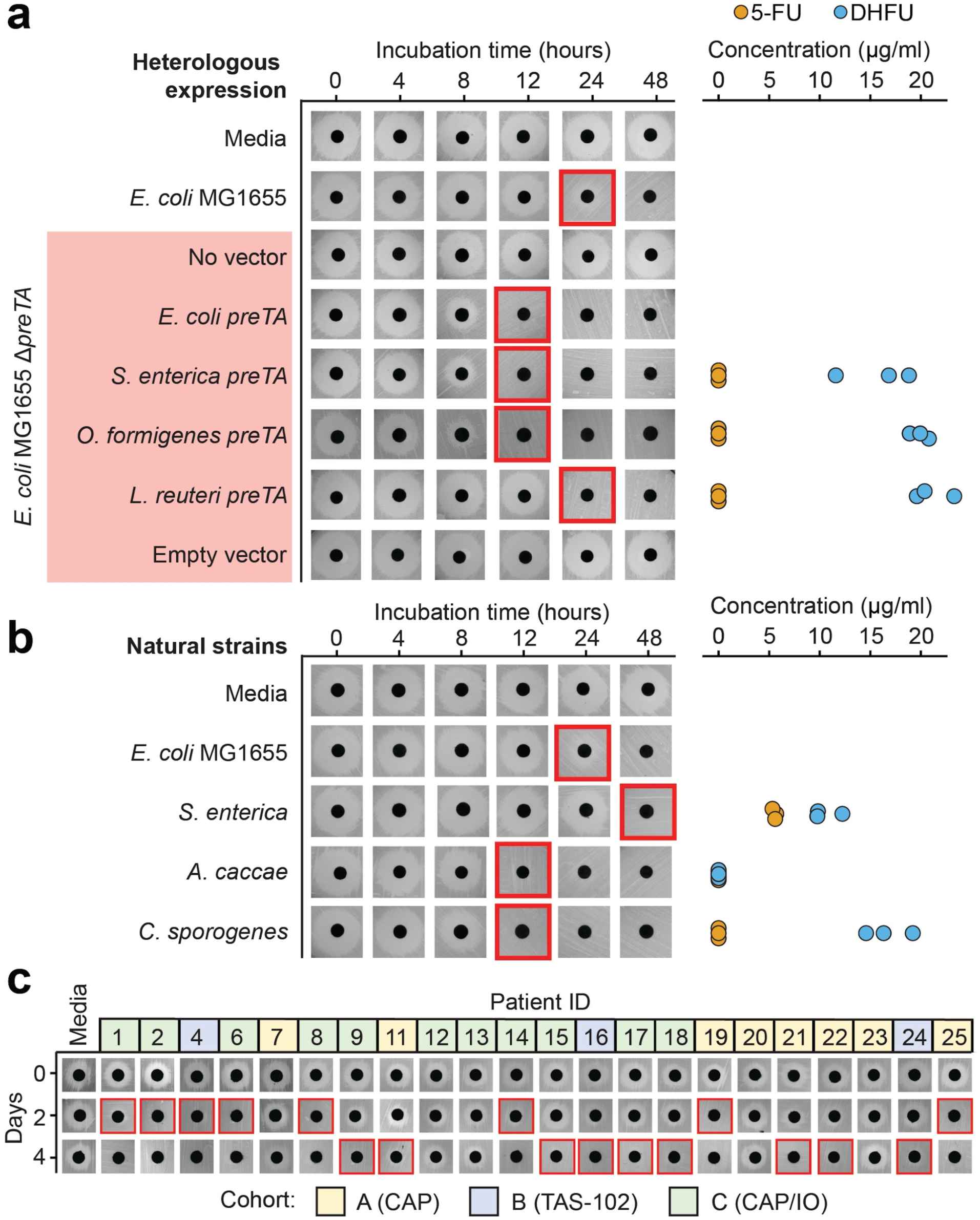
Bacterial *preTA* operons from diverse strains, *preTA* encoding natural strains, and pooled isolates from patient samples are capable of 5-FU inactivation. **(a)** *preTA* operons from *Salmonella enterica* LT2 (DSM17058), *Oxalobacter formigenes* ATCC35274, and *Lactobacillus reuteri* DSM20016 were amplified and integrated into the chromosome of *E. coli* MG1655 Δ*preTA*. Strains were incubated with 20 μg/ml 5-FU and assayed for residual drug using disk diffusion (0-48 hours incubation) and LC-QTOF/MS (48 hours). *E. coli* MG1655 *preTA* complementation and empty vector controls are also included on the *ΔpreTA* background. **(b)** We also tested strains predicted to encode the *preTA* operon: *Salmonella enterica* LT2, *Anaerostipes caccae* DSM14662, and *Clostridium sporogenes* DSM795. Residual 5-FU was assayed as in panel a. Of note, we did not detect either compound from *A. caccae*, suggesting that this strain may further metabolize DHFU. **(c)** Bacterial isolates from 22 CRC patient stool samples in the GO Study (**Supplementary Table 12**) were isolated on McConkey agar, pooled, and incubated in BHI^+^ with 5-FU for 4 days. Residual 5-FU levels were assessed using a disk diffusion assay. Colors indicate the treatment cohort: (A) CAP (yellow); (B) TAS-102 (blue); and (C) CAP plus immunotherapy (green). Red boxes in panels a-c indicate the first timepoint without a clear zone of inhibition.

**Extended Data Figure 10.**
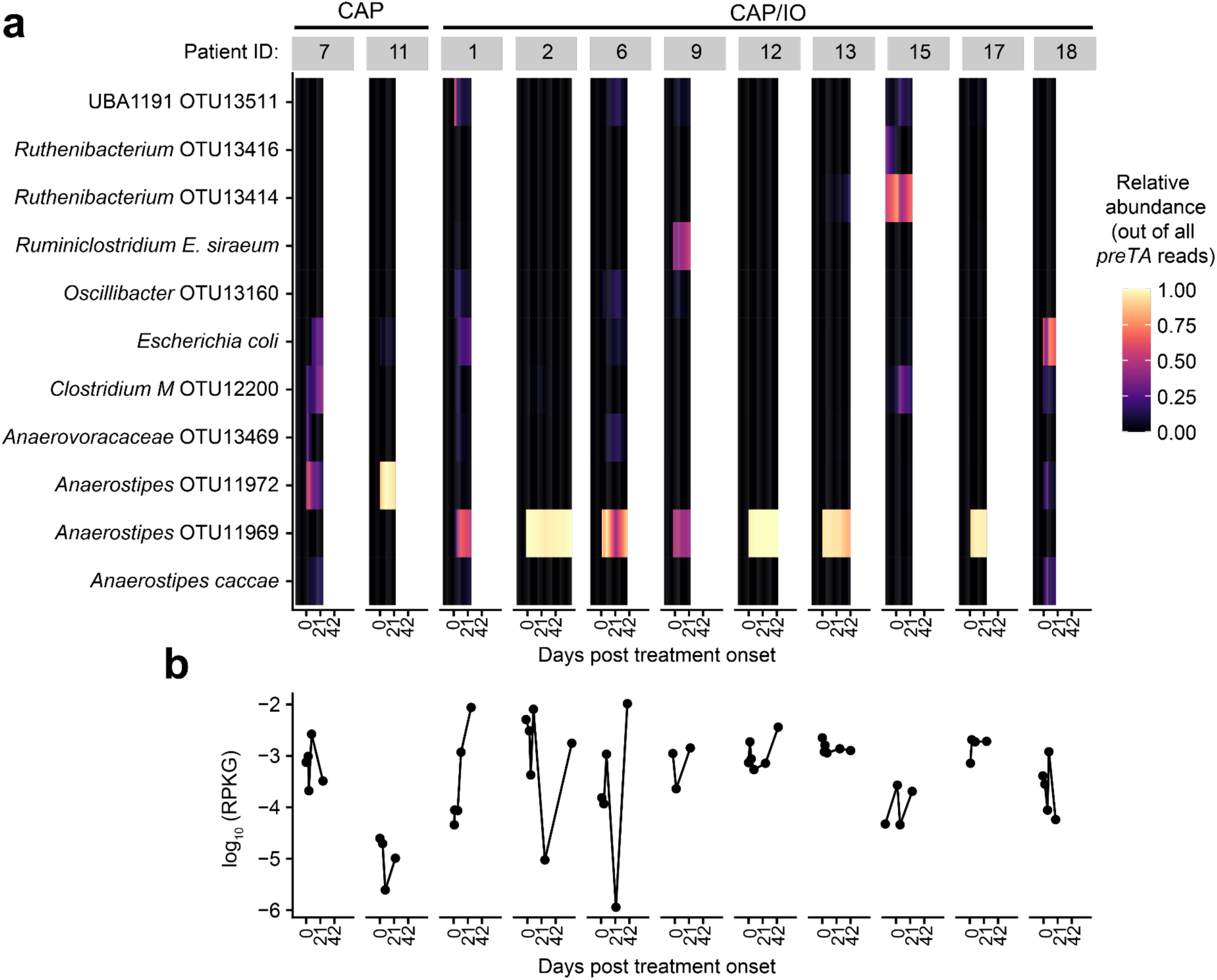
*preTA* sources and abundance in CRC patients during fluoropyrimidine treatment. **(a)** Fraction of total *preTA* reads mapping to different species in metagenomic samples from GO Study patients undergoing CAP treatment with or without pembrolizumab immunotherapy (IO). Heatmap values were linearly interpolated between samples (filled circles in panel **b**). **(b)** Total abundance, as log_10_ RPKG, of the *preTA* operon in the gut microbiome prior to and during treatment with the oral fluoropyrimidine CAP with or without IO, as shown in Fig. 4d. Lines connect measurements (filled circles) for the same patient. One zero RPKG value was replaced with half the minimum non-zero value prior to taking the logarithm. The first day of treatment is defined as day 1. Days are the same between panels.

