## Supplementary Information for "A conserved enzyme found in diverse human gut bacteria interferes with anticancer drug efficacy"

**SUPPLEMENTARY METHODS**

**Chemicals.** All materials were purchased from the specified supplier and used without further purification unless otherwise stated. 5-Fluorouracil (5-FU; ≥99% purity), 5-Fluorodeoxyuridine (FUDR; ≥99% purity), imidazol, and dithiothreitol were purchased from Millipore Sigma (St. Louis, MO). Capecitabine (CAP; ≥98% purity) used in this study was from Santa Cruz Biotechnology (Dallas, TX). Dihydrofluorouraucil (DHFU; ≥99% purity) was purchased from Toronto Research Chemicals (Toronto, Canada). Internal standards 5-FU $C^{13}$, $N_2^{15}$ (5-FU-IS; ≥99% purity) and 5-fluorodihydropyrimidine-2,4-dione-$C^{13}$, $N_2^{15}$ (DHFU-IS; ≥99% purity) were purchased from Santa Cruz Biotechnology Inc. (Santa Cruz, CA). Solvents used for sample preparation were methanol (MeOH), acetonitrile (ACN), and water and were purchased from Honeywell-Burdick & Jackson (Muskegon, MI). Benzamidine and 4-(2-aminoethyl)benzenesulfonyl fluoride hydrochloride (AEBSF) were purchased from RPI (Mt Prospect, IL). 2-mercaptoethanol was purchased from BioRad (Hercules, CA). Isopropyl β-D-1-thiogalactopyranoside (IPTG) and 4-(2-hydroxyethyl)piperazine-1-ethanesulfonic acid (HEPES) were purchased from Thermo Fisher Scientific (Waltham, MA).

**Bacterial strain collection.** Gut bacterial strains were obtained from the DSMZ (Braunschweig, Germany) and routinely cultured in $BHI^+$ media (**Supplementary Table 1**). Strains were mapped to the MIDAS v1.0 database[1]. If only one MIDAS species matched the strain species name, that MIDAS species was selected. If multiple MIDAS species matched, genomes in the MIDAS v1.0 database were filtered by whether the "culture collection" or "strain" metadata contained the DSMZ identifier, and then the most common remaining MIDAS species ID was kept. If no genomes matched the DSMZ identifier, the most common MIDAS species ID for that species name was selected. Abundance data came from a meta-analysis of five healthy cohorts described previously[2]. Abundances were summed across runs corresponding to the same individual, and then normalized per-individual to yield relative abundances for each species. The weighted mean of relative abundances was then taken per-species, with weights corresponding to the square root of the sample size. Four species were not represented in the MIDAS v1.0 database.

**Minimum inhibitory concentration (MIC) determinations.** All procedures and incubations were performed in an anaerobic chamber (COY Laboratory Products Inc.) with the following atmosphere: 10% $H_2$, 5% $CO_2$, and 85% $N_2$. Reagents were equilibrated in the anaerobic chamber for at least 24 hours before using. MIC determinations were performed in $BHI^+$ media: Brain Heart Infusion broth supplemented with L-cysteine hydrochloride (0.05% w/v), hemin (5 µg/ml), and vitamin K (1 µg/ml). 5-FU was dissolved in dimethyl sulfoxide (DMSO), supplemented at 1% (v/v) in $BHI^+$ media, and assayed at concentrations ranging from 0.1-1000 µg/ml. CAP was directly resuspended in $BHI^+$ media and assayed at concentrations ranging from 0.02-10 mg/ml. Each strain was inoculated in 5 ml of $BHI^+$ media in a Hungate tube and incubated at 37°C for 24-48 hours, depending on the time required to reach stationary phase. After reaching stationary phase, each strain was diluted to $OD_{600nm}$ of 0.1, further diluted 1:100, and 50 µl was used to inoculate a 96-well plate containing drug for a final volume of 100 µl. Plates were incubated at 37°C for 24-48 hours depending on the strain. The MIC was determined as the concentration of drug that inhibited the growth of a bacterial strain >90% relative to the growth control supplemented with vehicle. MIC assays were performed in duplicate. *Escherichia coli* BW25113 was used to perform the media effects and uracil rescue experiments. These MICs were conducted in triplicate in M9 minimal salts media (M9MM) provided by BD Difco (Franklin Lakes, NJ) supplemented with 0.4% glucose. Uracil from Acros Organics (Geel, Belgium) was supplemented at the indicated concentrations. Assays were performed in a 96-well plate with 100 µl total volume and incubated aerobically at 37°C for 24 hours. *E. coli* mutants deficient in various pyrimidine metabolism genes were obtained from the National BioResource Project (NBRP; Shizuoka, Japan) were assayed in duplicate in M9MM supplemented with 0.4% glucose and 5-FU or CAP.

**Transcriptional profiling (RNA-seq).** In triplicate, 100 ml of LB media was inoculated with an overnight culture of *E. coli* MG1655 at a ratio of 1:100. These cultures were incubated aerobically at 37°C with shaking at 250 rpm and anaerobically at 37°C without shaking. Two independent experiments were conducted for treatments under aerobic conditions. Cultures were grown to a mid-exponential phase ($OD_{600nm}$ of 0.5-0.6 and 0.26-0.28 under aerobic and anaerobic conditions respectively) and then 25 ml of culture was added to 25 ml of pre-incubated media with the following conditions: (1) fresh LB (control); (2) 5-FU (final concentration of 16 µg/ml); and (3) CAP (final concentration of 2.5 mg/ml). Drug concentrations were selected to represent 0.5X MIC under the selected media conditions. Cultures were incubated for an additional 30 min and then 4 ml of cells were harvested and immediately frozen. Bacterial pellets were resuspended with TRI reagent and then subject to mechanical lysis using MP Biomedicals Lysing Matrix E tubes (Solon, OH). Following extraction with chloroform and precipitation with ethanol, RNA was purified using the Life Technologies PureLink RNA Mini Kit (Carlsbad, CA). DNA was removed by using the Life

Technologies PureLink DNase Set (Carlsbad. CA). Depletion of rRNA was accomplished using the Illumina Bacterial Ribo-Zero rRNA Removal Kit (San Diego, CA; experiment 1) and Invitrogen Ribominus Bacterial Transcriptome Isolation Kit (experiment 2). RNA fragmentation, cDNA synthesis, and library preparation were performed using the NEBNext Ultra RNA library Prep Kit for Illumina and NEBNext Multiplex Oligos for Illumina (Dual Index Primers) (Ipswich, MA). Samples were dual-end sequenced (2 x 75 bp) using the MiSeq V3 platform (experiment 1) and NextSeq Mid Output platform (experiment 2; **Supplementary Table 2**). Reads were mapped to the *E. coli* MG1655 genome sequence (NCBI Reference Sequence: NC_000913.3) using Bowtie2[3] and HTSeq[4] was used to count the number of reads to *E. coli* genes. Differential gene expression was analyzed using DESeq[5]. Differentially expressed genes following drug treatment were defined as transcripts exhibiting an absolute log2 fold change ≥1 and a FDR<0.1 relative to the no drug control.

**Generation and analysis of 5-FU-resistant mutants.** For the generation of spontaneous 5-FU-resistant mutants on agar media, cultures of *E. coli* MG1655, *Bacteroides fragilis* DSM2151 and *Bacteroides ovatus* DSM1896 were grown anaerobically in $BHI^+$ liquid media overnight. The following day, 100 µl of cultures (~$1x10^9$ CFUs/ml) were plated on $BHI^+$ agar media supplemented with 1 mg/ml of 5-FU and incubated anaerobically for 48 hours. Representative 5-FU-resistant colonies were picked and restreaked on $BHI^+$ agar supplemented with 5-FU. For the generation of 5-FU-resistant mutants through passage in liquid media, an overnight culture was subcultured 1:100 into fresh $BHI^+$ liquid media with 50 µg/ml of 5-FU. Following overnight incubation, bacterial cultures were subcultured 1:100 into fresh $BHI^+$ liquid media with 250 µg/ml of 5-FU. Finally, a third 1:100 subculturing was performed in fresh $BHI^+$ liquid media supplemented with 1 mg/ml of 5-FU. Following overnight incubation, a bacterial suspension was streaked for single colonies on $BHI^+$ agar media supplemented with 1 mg/ml of 5-FU. 5-FU MICs were determined as described above. Genomic DNA was prepared using the DNeasy Blood and Tissue Kit (Qiagen, Hilden, Germany) according to the manufacturer's directions for Gram-negative bacteria. The *upp* gene, encoding uracil phosphoribsyltransferase, was amplified by PCR using primers upp_Ecoli_F and upp_Ecoli_R for *E. coli* and upp_Bfrag_F and Bfrag_R for *B. fragilis* (see **Supplementary Table 10** for primer list). Sanger sequencing of the PCR products was performed by GENEWIZ (South Plainsfield, NJ). For whole genome sequencing, genomic DNA was fragmented to 350 bp on a Covaris S2, and libraries prepared using a TruSeq DNA PCR-Free Library Prep Kit (Illumina) following the manufacturer's instructions. Samples were quantified using a Qubit dsDNA HS Assay, pooled, quantified by qPCR using a KAPA Library Quantification Kit (KAPA Biosystems), and submitted to the UCSF Institute for Human Genetics Genomics Core Facility for 100-base sequencing on an Illumina HiSeq platform. Reads were adapter trimmed using Trimmomatic[6] and mutations identified using Snippy[7] against reference genome sequences for *Bacteroides ovatus* ATCC 8483 (NZ_CP012938.1) and *Escherichia coli* K12 MG1655 (NC_000913.3).

**5-FU inactivation assay.** Starter cultures of gut bacterial strains were grown anaerobically in 5 ml of $BHI^+$ media in Hungate tubes at 37°C. The following day, bacterial cultures were diluted 1:100 into Hungate tubes with 5 ml of fresh $BHI^+$ media supplemented with 5-FU at 20 µg/ml. Over time, conditioned media samples were harvested, centrifuged to remove cells, and assayed for residual 5-FU levels. *E. coli* BW25113 Δ*yjjG*, which is a 5-FU-hypersensitive strain[8], was obtained from National BioResource Project (NBRP, Shizuoka, Japan) and used as the indicator organism for the disk diffusion assay. An overnight culture of *E. coli* BW25113 Δ*yjjG* was diluted to an $OD_{600nm}$ of 0.1 in saline. Using sterile swabs, the bacterial suspension was used to inoculate M9MM plus 0.4% glucose and 0.2% casamino acids agar plates and a paper disk was overlaid. Residual 5-FU concentration was assayed by diluting conditioned media samples 1 in 5 in water and 1 µl was applied on the paper disk. Agar plates were incubated aerobically overnight at 37°C. Under these conditions, our assay allowed the detection of between 1-20 µg/ml of 5-FU. Strain designations for bacteria carrying the *preTA* operon that were tested for inactivation but not included in our initial strain collection are: *Anaerostipes caccae* DSM14662 and *Clostridium sporogenes* ATCC15579.

**Liquid chromatography-mass spectrometry analysis of 5-FU and metabolites.** We used two separate high-resolution mass spectrometry instruments based upon staff and instrument time constraints. The two protocols are described in detail below.

Protocol A: 5-FU and DHFU from conditioned media samples were analyzed by liquid chromatography-quadrupole time-of-flight mass spectrometry (LC-QTOF/MS) using an Agilent LC 1260-QTOF/MS 6550 instrument (Santa Clara, CA). Conditioned media samples (25 µl) were spiked with 2.5 µl of a MeOH mix containing internal standards (10 µg/ml 5-FU $^{13}C^{15}N_2$, and 100 µg/ml DHFU $^{13}C^{15}N_2$). Protein from conditioned media samples were precipitated with an equal volume of MeOH. The resulting extracts were dried down and thereafter reconstituted with 25 µl of a 10% ACN in water solution and further diluted 20X prior to LC-QTOF/MS analysis. Chromatographic separation was achieved using a Phenomenex Luna $NH_{-2}$ (50 x 2 mm, 3 mm) connected to a Phenomenex Securityguard™ guard cartridge (4x2 mm) (Torrance, CA) at 35°C.

The mobile phase consisted of 20 mM ammonium hydroxide, 20 mM ammonium acetate, 5% ACN in water as solvent A, and 100% ACN as solvent B. A flow rate of 0.8 ml/min was used at the following gradient elution profile: 90% B at 0-0.5 min, gradient to 30% B from 0.5-2 min, 90% B at 2-5 min. 2.5 µl of clarified conditioned media was injected for LC-QTOF/MS analysis. The autosampler was maintained at an internal temperature of 4°C. Eluate from the column was ionized in the QTOF/MS using an electrospray ionization source (ESI) in negative polarity. Qualitative confirmation of 5-FU and DHFU in each sample was done using the Agilent MassHunter Qualitative Analysis (Santa Clara, CA) software. The criteria used for confirmation were as follows: mass error ≤ 10 ppm; retention time within 0.15 min; and target score ≥ 75 (overall indication of match based on mass error, retention time match and isotope abundance match). Quantitative analysis for both 5-FU and DHFU was performed using an isotope dilution method with an 11-point calibration curve run in triplicate. Data analysis was performed using Agilent MassHunter Quantitative Analysis software. The linear regression of the peak area ratio was weighted 1/x for both 5-FU and DHFU with respective linear regression coefficients of $R^2$=0.996 and $R^2$=0.992. The lower limits of quantitation (LLOQ) for 5-FU and DHFU were 100 ng/ml and 4 µg/ml, while the observed upper limits of quantitation (ULOQ) were 1.25 µg/ml and 30 µg/ml, respectively.

Protocol B: CAP, 5-FU, and DHFU from conditioned media samples were analyzed by liquid chromatography-triple quadrupole mass spectrometry (LC-MS/MS) using a SCIEX Triple Quad 7500 instrument with a linear ion QTRAP (Redwood City, CA). Conditioned media samples (10 µl) were spiked with 1 µl of a MeOH mix containing internal standards (10 µg/ml 5-FU $^{13}C^{15}N_2$, 100 µg/ml DHFU $^{13}C^{15}N_2$, and 10 µg capecitabine-$^2H_{11}$). Protein from conditioned media samples were precipitated with 60 µL of MeOH. The resulting extracts were dried down and thereafter reconstituted with 500 µl of a 10% ACN in water solution. Chromatographic separation was achieved using a Phenomenex Synergi column 4 µM Fusion RP-80 (50 x 2 mm) at 35°C. The mobile phase consisted of methanol + 0.1% formic acid for solvent A and HPLC-grade water + 0.1% formic acid as solvent B. A flow rate of 0.4 ml/min was used at the following gradient elution profile: 0% B at 0-2 min, gradient to 100% B from 2-5.9 min, gradient to 0% B at 5.9-6 min. The autosampler was maintained at an internal temperature of 4°C. Eluate from the column was ionized in the LC-MS/MS using an electrospray ionization source (ESI) in positive polarity. Quantitative analysis for both 5-FU and DHFU is performed using an isotope dilution method with a 9-point calibration curve run in duplicate. Data analysis was performed using the built-in SCIEX OS Software. The linear regression of the peak area ratio was weighted 1/x for CAP with a linear regression coefficient of $R^2$ = 0.995. The lower limits of quantification (LLOQ) for CAP was 1 nM and the upper limit of quantification (ULOQ) for CAP was 2.5 µM. The quartic regression of the peak area ratio was weighted 1/x for both 5-FU and DHFU with quartic regression coefficients of $R^2$=0.998 and $R^2$=0.987, respectively. The lower limits of quantitation (LLOQ) for 5-FU and DHFU were 1 µM and 31 µM, while the observed upper limits of quantitation (ULOQ) were 250 µM for both.

**Liquid chromatography-mass spectrometry analysis of 5-FU from plasma samples**
Protocol A: The validated LC-QTOF/MS method for the conditioned media described above was adopted to measure 5-FU in the plasma samples. Mouse plasma (9-18 µl) was extracted using protein precipitation with methanol at 3:1 methanol: plasma volume ratio Similarly, the resulting extracts were dried down and thereafter reconstituted in 10% acetonitrile (ACN) at the same volume as the plasma sample. The final extract obtained was then run using the same qualitative and quantitative analyses applied to the conditioned media. The LLOQ and ULOQ observed for 5-FU in mouse plasma were 20 ng/ml and 1500 ng/ml, respectively.

Protocol B: The validated LC-MS/MS method for conditioned media was adapted to measure 5-FU in plasma. Mouse plasma (10 µl) was extracted by protein precipitation with methanol (60 µl). The extract was dried under a gentle stream of nitrogen gas and reconstituted in 30 µl of 10% acetonitrile. The final extract was then run using the same quantitative analyses applied to the conditioned media. The LLOQ and ULOQ observed for 5-FU in mouse plasma was 5 ng/ml and 1500 ng/ml, respectively.

**Bacterial genetics.** Markerless mutant strains of *E. coli* BW25113 and *E. coli* MG1655 devoid of the *preTA* operon were constructed using the recombineering protocol[9] with the pSIJ8 vector as previously described[10]. The *preTA* operon was replaced using allelic exchange by electroporation of a PCR product containing the kanamycin cassette[11,12], which was generated using the primers preTA-P1-KEIO_F and preTA-P1-KEIO_R (**Supplementary Table 10**) and using the *E. coli* BW25113 Δ*yjjG* genomic DNA as a template. Allelic exchange of the *preTA* operon with the kanamycin cassette was confirmed by PCR. *preTA* operons identified in diverse gut bacterial genomes were amplified and cloned into the pINT1 vector[13] using primers listed in **Supplementary Table 10**. A list of *preTA* operons including sequences used for heterologous expression experiments can be found in **Supplementary Table 10**. The *preTA* operon from *O. formigenes*, *preTA-Of*, was codon optimized for expression in *E. coli* and synthesized by

GENEWIZ (South Plainfield, NJ). Integration of the *preTA*-pINT1 constructs into *E. coli* MG1655 Δ*preTA* was performed as previously described[10,13]. Engineered strains were made streptomycin resistant by using the recombineering protocol[9] using pSIJ8[10], as described above. A spontaneous streptomycin-resistant mutant of *E. coli* MG1655 was created by plating an overnight culture on LB agar supplemented with 100 µg/ml of streptomycin and incubated at 37°C overnight. Streptomycin-resistant colonies were picked and restreaked on LB agar supplemented with 100 µg/ml of streptomycin to confirm their resistant phenotype. The *rpsL* gene from streptomycin-resistant isolates were amplified by PCR using the primers rpsL_F (5'-CGTGGCATGGAAATACTCCG-3') and rpsL_R (5'-GCATCGCCCTAAAATTCGGC-3'). PCR products were sequenced by GENEWIZ (South Plainsfield, NJ) using Sanger sequencing with primers rpsL_F and rpsL_R. We selected the RpsL K42R mutation for future experiments[14]. Mutant *rpsL* PCR product was transferred to *preTA* operon engineered strains described above using the recombineering protocol[9] with the pSIJ8 vector as previously described[10] and selected by plating on LB agar supplemented with 100 µg/ml streptomycin. Streptomycin-resistant colonies were picked and restreaked to confirm resistance phenotype. The *rpsL* gene from these colonies were also amplified by PCR and sequenced to confirm point mutation.

**Cancer cell culture assays.** HCT-116 cells (ATCC CCL-247) were acquired from the ATCC (Manassas, VA) and grown in McCoy's 5A medium supplemented with 10% fetal bovine serum (FBS) and Gibco antibiotic-antimycotic solution. Cells were grown at 37°C with 5% $CO_2$. Approximately 2.4 x $10^4$ HCT-116 cells were seeded into a Corning Costar TC-treated 48-well plate in a total volume of 250 ml culture media. Following 24 hours, 28 µl of conditioned media was added to HCT-116 cells to assay for residual 5-FU activity. Conditioned media samples from *E. coli* MG155 cultures and related *preTA* constructs were grown anaerobically for 3 days in $BHI^+$ media with 5-FU at a concentration of 20 µg/ml with a final DMSO concentration of 0.05%. Conditioned media samples were sterilized by passage through a 0.2 µm filter prior to assaying against HCT-116 cells. Cell proliferation was measured using the Promega CellTiter Non-Radioactive Cell Proliferation Assay (Madison, WI). Cell proliferation was normalized to HCT-116 cells that were supplemented with water instead of conditioned or sterile culture media.

**Cloning, expression, and purification of *E. coli* PreTA.** Polycistronic pCOLADuet-1 constructs containing $His_6$_*Ec*PreT and untagged *Ec*PreA and were kindly provided by Professor Catherine Drennan's Lab at MIT. pDB1282 was kindly provided by Professor Dennis Dean at Virginia Tech. Polymerase Chain Reaction followed by Gibson Assembly was used to remove the N-terminal

*Ec*PreT $His_6$ tag and to include a C-terminal tobacco etch virus (TEV) protease site followed by a $His_6$-tag to *Ec*PreA such that the final expression vector was pCOLADuet-1_*Ec*PreT_*Ec*PreA-TEV-$His_6$ (see **Supplementary Table 10** for primer list). The final expression construct was confirmed by Sanger Sequencing. pCOLADuet-1_*Ec*PreT_*Ec*PreA-TEV-$His_6$ was co-transformed with pDB1282 into *E. coli* Rosetta2 (DE3) cells and selected for with kanamycin, chloramphenicol, and ampicillin. pDB1282 ($Amp^R$) contains the iron sulfur cluster (*isc*) biosynthetic operon from *Azotobacter vinelandii* under the control of the *E. coli* arabinose promoter $P_{ara}$ which has been shown to increase the incorporation of Fe-S clusters during heterologous expression[15] .

A single colony of *E. coli* Rosetta2 (DE3) harboring the expression construct and pDB1282 was inoculated in TB media (10 g/L NaCl) supplemented with 50 µg/mL kanamycin, 25 µg/mL chloramphenicol, and 100 µg/mL ampicillin and grown overnight at 37 °C with shaking at 225 rpm. The overnight culture was diluted 1:100 into 1 L of TB media supplemented with the same antibiotics as before and grown at 37 °C with shaking at 225 rpm until $OD_{600nm}$ = 0.6. Arabinose was added to a final concentration of 0.2% (w/v) and the cultures were allowed to continue growing until $OD_{600nm}$ > 2.0. At this point, the cultures were cooled in an ice bath for 30 min and the following was added to the cultures: 10 µM uracil, 20 µM riboflavin, 10 µM cysteine, 10 µM ferric chloride, and 500 µM IPTG. The temperature of the incubator shaker was lowered to 18 °C and the protein was allowed to express with shaking at 225 rpm overnight. Cell pellets were harvested by spinning down 1 L cultures in a pelleting centrifuge, collected in tubes, snap frozen in liquid nitrogen, and stored at –80 °C.

All following purification steps were carried out at 4 °C. Cell pellets were thawed in an ice water bath and then resuspended by vortexing in 150 mL of Lysis Buffer that consisted of Buffer A (50 mM $K_2HPO_4$ pH 8.0, 200 mM NaCl, 5% glycerol, 0.22 µm filtered) supplemented with 25 mM benzamidine, 1 mM AEBSF, 0.01 mg/mL bovine DNase, 0.1 mg/mL lysozyme, and 5 mM 2-mercaptoethanol. Cells were lysed by sonication using a QSonica Q500 Sonicator at 30% amplitude for 5 minutes of total process time (2 seconds on, 4 seconds off). Whole cell lysate was clarified by centrifuging at 30,000 *g* for 30 minutes. Clontech TALON Superflow resin (3 mL bed volume) was equilibrated with 10 column volumes (CV) of lysis buffer and then added to the supernatant of the clarified lysate and rocked for 30 minutes. The supernatant and resin were applied to a gravity column which was allowed to flow until the resin was collected. Next, the column was washed with 10 CV of Buffer A supplemented with 5 mM 2-mercaptoethanol and steps of increasing imidazole concentration until the protein was eluted off with Buffer B (50 mM $K_2HPO_4$ pH 8.0, 200 mM NaCl, 150 mM imidazole, 5% glycerol, 0.22 µm filtered). The elution fractions were dialyzed in a ratio of sample to dialysate of 1:500 for 2 hours against Buffer C (50 mM Tris pH 7.5, 150 mM NaCl, 1 mM 2-mercaptoethanol) prior to the addition of TEV protease, and the dialysis was allowed to continue overnight. The cleaved protein was collected in the flowthrough of a TALON Superflow subtractive column and then spin concentrated to 0.5 mL. Finally, the protein was injected onto a GE Superdex 200 Increase 10/300 GL size exclusion column that was pre-equilibrated with Buffer D (50 mM Hepes pH 7.4, 150 mM NaCl, 5 mM DTT, 5% glycerol, 0.22 µm filtered). Purity was assessed by SDS-PAGE and cofactor incorporation was evaluated with UV-Visible absorption spectroscopy ($A_{378}/A_{280} > 0.35$)[16]. The main band was pooled and concentrated to a concentration of 3-5 mg/mL as assessed by Nanodrop $A_{280}$ (ε = 114.6 $M^{-1}cm^{-1}$, MW = 178.74 kDa), snap frozen in liquid nitrogen, and stored at -80 °C.

**Biochemical characterization of *E. coli* PreTA.** To collect UV-Visible absorption spectra in different oxidation states, *Ec* PreTA was brought into an anaerobic chamber and $Na_2S_2O_4$ was added in an approximately equimolar amount. Samples were placed in a septum-sealed 1 cm path-length quartz cuvette inside the glovebag, and UV−Visible absorption spectra were recorded on a Cary 3500 spectrophotometer (Agilent Technologies). Analytical Size Exclusion Chromatography was performed by injecting three subsequent samples onto a GE Superdex 200 Increase 10/300 column pre equilibrated with Buffer D: 0.25 mg/mL blue dextran for void volume determination, 0.25 mg/mL Protein Standard Mix 15-600 kDa (Millipore Sigma), and a 2 mg/mL *Ec* PreTA sample. Samples were run at 0.5 mL/min. The standard curve was generated by plotting $\log_{10}$ of the molecular weight of the protein standard in Da versus the fractional elution volume ($V_{elution}/V_{void}$). Enzyme assays to confirm product formation were performed by bringing the necessary reagents into a COY anaerobic chamber and diluting into anaerobic Buffer E (25 mM Hepes pH 7.4, 25 mM NaCl, 0.22 µm filtered). Substrates were prepared at only twice the final concentration to ensure complete solubility. The final reaction contained Buffer E supplemented with 5 mM DTT, 800 µM NADH, 400 µM substrate, and was initiated by addition of 100 nM *Ec* PreTA at 25 °C. Aliquots were quenched into 2% formic acid. Reactions were analyzed by reverse-phase HPLC to observe product formation by running an isocratic method of 100% Buffer F (0.1% formic acid (FA) in $dH_2O$, 0.22 µM filtered, degased) followed by a cleaning cycle of 100% Buffer G (0.1% formic in acetonitrile, degased) and reequilibration in Buffer F over a Phenomenex Kinetex 150 x 4.6 mm, 2.6 µM C18 column. Reaction product masses were confirmed by liquid chromatography high resolution mass spectrometry (LC-HRMS) on a SCIEX TripleTOF 6600+. Reactions were applied to a Phenomenex Kinetex F5 (2.6 uM, 100A, 150 x 2.1 mm) column and products were separated with a gradient of 100% water (+ 0.1% FA) to 95% acetonitrile (+0.1% FA) over 12 minutes. Independent data acquisition (IDA) of TOF MS (DP 30, CE 10, ISG1 60,

ISG2 80, CUR 35, TEM 400, ISVF 5500) and product ion (DP 50, CE 35, CES 15, IRD 66, IRW 24) spectra were obtained in positive ionization mode. To measure the steady-state kinetics of *Ec* PreTA, depletion of NADH was measured by following absorbance decreases at 340 nm throughout the course of the reaction. The final reaction contained Buffer G (100 mM $K_2HPO_4$ pH 7.4) supplemented with 5 mM DTT, 400 μM NADH, variable concentrations of substrate, and were initiated by addition of 10 nM *Ec* PreTA for the forward reactions at 25 °C and Buffer G (100 mM $K_2HPO_4$ pH 7.4) supplemented with 5 mM DTT, 1600 μM $NAD^+$, variable concentrations of substrate, and were initiated by addition of 100 nM *Ec* PreTA for the reverse reactions at 25 °C. Initial rates were normalized to a control reaction where no substrate was added and calculated from the linear phase of the time course during which 5-10% of the NADH substrate was consumed.

**CAP pharmacokinetic studies.** All animal experiments were approved by the University of California, San Francisco (UCSF) IACUC. In the gnotobiotic pharmacokinetics experiment, 8-10 week-old female germ-free BALB/c mice were mono-colonized with isogenic *E. coli* strains with various *preTA* operon statuses (Δ*preTA, wt*, *preTA*$^{++}$*).* 7 days post-colonization, the mice were fasted overnight prior to drug administration. CAP used in animals was from LC Laboratories (Woburn, MA) and was freshly prepared in 40 mM citrate buffer (pH 6.0) containing 5% (w/v) gum arabic. CAP was delivered by oral gavage (0.2 ml) at a dose of 500 mg/kg. Blood samples (~25 μl per time point) were obtained from the mouse tail vein at 0, 0.5, 1, 2, 3, 4, 6, 8 hours post CAP administration. Blood samples were collected in Fisherbrand heparinized glass microhematocrit capillary tubes (Waltham, MA) and centrifuged at 3500×g for 15 min to recover blood plasma, which was stored at -80°C until analysis.

In the conventionally-raised (CONV-R) specific pathogen free pharmacokinetic experiments, 8-10 week-old male BALB/c mice (Taconic Biosciences, Model#: BALB-M) were colonized with either streptomycin-resistant *ΔpreTA* or *preTA*$^{++}$ *E. coli* MG1655 isogenic strains using the streptomycin mouse model (as described in the CAP xenograft tumor model below). They were colonized for 1 week prior to pharmacokinetic experiments. After the overnight fast, CAP was delivered by oral gavage (0.2ml) at the dose of 500 or 1100 mg/kg. Blood was collected at 0, 0.5, 1, 1.5, 2, 3, and 5 hours post CAP administration and processed in the same procedure as described above for gnotobiotic pharmacokinetic experiment.

**CAP tumor xenograft model.** All animal experiments were approved by the University of California, San Francisco (UCSF) IACUC. Three independent experiments (see **Extended Data**

**Figs. 6a, 6g, and Fig. 3a** for designs of experiment 1, 2, and 3, respectively) were performed with female athymic nude mice at 6 weeks of age (Taconic Biosciences, model #: NCRNU-F). HCT-116 cells were grown in McCoy's 5A medium supplemented with 10% fetal bovine serum and 1% penicillin-streptomycin in 3x100 mm tissue-culture treated dishes at 37°C with 5% $CO_2$ to 85% confluence. Cells were detached with 2 ml of 0.05% trypsin and quenched with 10 ml of media. Cells were pelleted by centrifugation (200 g for 5 min), media removed, resuspended in 3.5 ml of PBS, and kept on ice for the remainder of the procedure. Cold cells were mixed with BD Matrigel™ Basement Membrane Matrix (Franklin lakes, NJ) at a 1:1 ratio. Each animal was put under isoflurane anesthesia, given artificial tears ointment, and received 100 µl of cell-Matrigel™ mix ($1x10^6$ cells) with a 27G1/2 needle in the subcutaneous space on the right flank. Tumors were grown for 19 days and 13 days before bacterial colonization and CAP treatment initiation in experiment 1 and 2, respectively. In experiment 3, bacterial colonization and CAP treatment were initiated when the tumors reached between 80-120 $mm^3$ on a rolling enrollment basis.

Mice were colonized with the engineered streptomycin-resistant *E. coli* strains described above (*E. coli* MG1655 Δ*preTA* and *E. coli* MG1655 preTA++) using the streptomycin mouse model[17], which we confirmed leads to stable colonization of *E. coli* at high abundance (*data not shown*). The day before colonization, mice were put on filtered streptomycin (5 g/L for experiment 1 and 3, 5 mg/L for experiment 2) water. Overnight cultures of engineered *E. coli* strains were pelleted by centrifugation, washed with an equal volume of sterile 0.85% saline, pelleted by centrifugation again, and resuspended in 1:10 sterile saline. Each mouse was colonized with 200 µl of this bacterial suspension by gavage. Mice were housed (n=4-5 mice/cage) according to colonization status and drug or vehicle treatments and were on streptomycin tap water for the duration of the tumor xenograft experiment. Streptomycin tap water was freshly prepared and replenished weekly. *E. coli* colonization levels were determined by culturing from fecal pellets. Freshly collected fecal pellets (1-2 pellets per animal) were weighed in a microcentrifuge tube and 1 ml of 1% (w/v) Bacto tryptone solution was added to each tube. Fecal pellets were resuspended by rigorously vortexing for 5 min and large particles were removed by allowing the suspension to settle by gravity for 5 min. 10-fold serial dilutions of fecal pellet suspensions were performed in 1% Bacto tryptone solution and 100 µl of these were spread on MacConkey agar supplemented with 100 µg/ml of streptomycin. Culture plates were incubated aerobically 37°C overnight. Colony-forming units (CFUs) per gram fecal sample was calculated by dividing CFUs by 0.1 ml (volume plated), dilution, and by sample weight. Colonization levels were monitored before and after CAP treatment.

CAP used in animals was from LC Laboratories (Woburn, MA) and was freshly prepared daily in 40 mM citrate buffer (pH 6.0) containing 5% (w/v) gum arabic. CAP was delivered by oral gavage (0.2 ml) at a dose of 100 mg/kg for a total of 15 doses over 17 days in experiments 1 and 3 and 18 doses over 22 days in experiment 2. Tumor dimensions were measured with a digital caliper 2-3 times a week. Tumor volume was calculated with the following formula: volume ($mm^3$) = (length × $width^2$)/2[18]. The remainder of the mice were allowed to reach the humane endpoint defined as tumor length >20 mm, tumor ulceration, and body condition score of ≤2.

**16S rRNA gene sequencing of fecal pellets from CAP tumor xenograft model**. Mouse fecal pellets were collected throughout the tumor xenograft experiment and stored at -80°C. DNA was extracted using a ZymoBIOMICS 96 MagBead DNA Kit (Zymo D4302) and 16S rRNA amplicon library was constructed[19] using dual error-correcting barcodes. Briefly, primary PCR was performed as a quantitative PCR using KAPA HiFi Hot Start kit (KAPA KK2502) and V4 515F/806R Nextera primers. The amplified products were diluted 1:100 in UltraPure DNase/RNase-free water and were indexed using sample-specific dual indexing primers. The reactions were quantified using Quant-iT PicoGreen dsDNA Assay Kit (Invitrogen P11496) and pooled at equimolar concentrations. The pooled library was quantified by qPCR using KAPA Library Quantification Kit for Illumina Platforms (KAPA KK4824) and sequenced on Illumina MiSeq platform. The demultiplexed sequences were processed using a 16S rRNA gene analysis pipeline (https://github.com/turnbaughlab/AmpliconSeq) and analyzed using qiime2R (https://github.com/jbisanz/qiime2R), phyloseq[20], and DESeq[5] in R. Briefly, the ASV tables, phylogenetic tree, taxonomy files from qiime2R outputs along with the sample metadata were used to build a phyloseq object using the phyloseq package. To control for uneven sequencing depth, the samples were sub-sampled to 10,000 sequencing reads. Alpha diversity, beta diversity, and community composition were analyzed using the phyloseq built-in functionalities. DESeq package was used to determine the differentially abundant taxa between the treatment groups.

**Bioinformatic analysis of the *preTA* operon in genomes and microbiomes.** Amino acid sequences of PreT and PreA from *Escherichia coli* (Genbank protein ID: AVI56642.1 and AVI56643.1), *Salmonella enterica* (AUX97235.1 and AUX97234.1), *Anaerostipes caccae* (EDR96331.1 and EDR96330.1), *Clostridium sporogenes* (EDU36024.1 and EDU36025.1), and *Oxalobacter formigenes* (ARQ46500.1 and ARQ46501.1), as well as the human DPYD protein (NP_000101.2), were downloaded from the NCBI Protein database[21]. T-COFFEE[22,23] was used to construct multiple alignments of PreT and PreA, each containing the five bacterial PreTA sequences plus human DPYD. Non-aligning columns were removed with trimAl[24], using "automated1" parameter selection. Using HMMER3[25], amino acid profile hidden Markov models (HMMs) were built from the trimmed alignments, and these HMMs were used to search against protein sequences from 9,411 complete genomes in RefSeq[21] (9,032 of which were bacterial). Significant (*e*-value < $1e^{-10}$) hits (5,622 to the PreA HMM and 12,849 to the PreT HMM) were filtered using RefSeq GFF3-formatted genome annotations to find PreT and PreA coding sequences that were from adjacent genes on the same strand in a bacterial genome (yielding 1,704 putative operons). Previously, gut prevalences[2] were calculated for species in the MIDAS v1.0 database[1]. To link our results with these, we first used the tool BURST to look across the RefSeq genomes for close matches to the 15 PhyEco marker genes[26] that MIDAS uses to perform taxonomic identification. We then matched genomes to MIDAS species by requiring at least 10 hits to the same MIDAS species above a 95% nucleotide identity threshold. Finally, to find putative *preTA* operons from uncultured bacterial strains, we analyzed a collection of draft metagenome-assembled genomes (MAGs; N=60,664 in total), each assembled from a single gut microbiome and representing 2,962 distinct gut OTUs in total[27], were searched in the same way as the RefSeq genomes. The phylogenetic species trees used for visualization were both constructed using IQ-TREE[28] on a concatenated protein alignment of single-copy marker genes. Specifically, for the tree of RefSeq genomes, we used HMMER3 and the PhyEco HMMs to identify the 15 single-copy universal marker genes selected in MIDAS, built alignments using Clustal Omega[29], trimmed columns with <60% occupancy, and finally built an approximate-maximum-likelihood tree from the concatenated alignment using IQ-TREE . Two *Methanobrevibacter smithii* genomes were incorporated into the alignment and used as an outgroup to root the tree. Certain Proteobacterial genera of mainly obligate endosymbionts (*Buchnera* and the *Candidatus* genera *Tremblaya, Nasuia, Carsonella, Portiera, Sulcia, Riesia,* and *Babela*) had very long branch lengths and were dropped from the tree to aid visualization. Also, so that species with hundreds of genomes did not dominate the visualization, when there were multiple genomes that had exactly the same species-level taxonomic annotation (both using the RefSeq NCBI annotations and using the MIDAS mapping) and the same prevalence estimates, a single genome was retained arbitrarily. The tree of MAGs was previously described[1,27]. To find orthologs of the *preTA* operon from our collection of gut bacteria used for screening, we used the *E. coli preTA* operon as the query in a tBLASTn[30] search of our specific organism list (**Supplementary Table 1**), which is publicly available.

To quantify *preTA* abundance in metagenomes without being misled by other *preT* or *preA* homologs, we applied a conservative five-step pipeline as follows. First, we built a database of genomic regions around the putative *preTA* operons that we identified from MAGs and Refseq genomes above. These regions included any intergenic sequence between the operon and the next annotated genes (or, for MAGs, the end of the contig, whichever was shorter). Second, we then extended this database by adding “decoy” regions, which included the coding and intergenic sequences around *preT* or *preA* homologs that matched our profile HMMs above but were not clustered into an operon. Third, we used vsearch[31] to align shotgun metagenomic reads to the database, retaining only reads that aligned to a putative operon and did not align better to a decoy. To reduce compute time while still allowing us to determine the probable origin of preTA reads, we ran the search in two stages: a) we initially reduced the database size using vsearch, clustering regions at 95% nucleotide identity, and then aligned metagenomes to this reduced database at a 95% ID threshold; then, b) we realigned any reads matching the reduced database against the full database, using a more stringent 97% ID threshold. Fourth, we added an additional coverage filter at the level of regions, dropping any reads mapping to regions that had highly uneven or partial coverage. Regions were kept if, across all samples in the dataset, a) both the preA and preT ORFs had non-zero coverage, and b) the median per-nucleotide coverage was >50% of the mean coverage. Fifth, the resulting read counts were normalized for average genome size using MicrobeCensus[32]. The pipeline was applied to two datasets: a previously published colorectal cancer (CRC) case-control study[33] and our own longitudinal study. The CRC dataset included 575 samples from 179 individuals, 141 of which were from a French population (88 controls and 53 with CRC) and the remaining 38 of which were from a German cohort (all with CRC). We followed the original authors’ classification of samples, dropping large adenomas and classifying small adenomas as controls. The longitudinal study had 54 samples from 11 CRC patients treated either with CAP alone (n=2) or CAP plus immunotherapy (n=9). For the taxonomic classification of preTA reads, we used the GTDB taxonomy[34] annotation of the best hit in our region database. Full source code can be found at https://bitbucket.org/pbradz/preta.

**Gut Microbiome and Oral Fluoropyrimidine Study in Patients with Colorectal Cancer (GO) sample collection and metagenomic sequencing.** This ongoing study is registered at ClinicalTrials.gov under the identifier NCT04054908, which includes detailed descriptions for the study population and enrollment criterias. All subjects provided informed consent to participate in the study, which was approved by the University of California, San Francisco (UCSF) Institutional Review Board. Patients with CRC who met the following criteria were recruited and screened for eligibility: (i) male or female aged 18 or older, (ii) histologically confirmed colorectal adenocarcinoma, (iii) expected to receive oral fluoropyrimidine therapy, (iv) able to read and speak English, and (v) willing and able to provide informed consent. Additionally, patients who met any of the following criteria were excluded: (i) known HIV positive diagnosis, (ii) prior chemotherapy, biologic or immunotherapy in the previous 2 weeks, (iii) concurrent rectal radiation therapy, (iv) exposure to antibiotics longer than 2 weeks in the last 6 months, or (v) exposure to antibiotics in the past 4 weeks prior to starting oral chemotherapy. Patients were enrolled to one of the following cohorts: Cohort A received oral CAP treatment as part of standard-of-care therapy; cohort B received TAS-102 (trifluridine-tipiracil) with or without Y-90 radioembolization; and cohort C received CAP plus immunotherapy (pembrolizumab) and bevacizumab as part of a clinical trial. The enrollment criteria were amended to enroll patients in cohort A who received concurrent rectal radiation therapy with CAP (see **Supplementary Table 12** which includes one patient who received concurrent rectal radiation therapy). CAP and TAS-102 tablets were taken per the FDA labels for oral dosing. CAP tablets were taken twice daily on days 1-14 of a 21 day cycle; TAS-102 tablets were taken twice daily on days 1-5 and 8-12 of a 28 day cycle. Stool collection occurred at baseline before chemotherapy initiation and on Day 1 of Cycles 1, 2 and 3. During cycle 1, stools were collected at two additional time points: (i) 2 days after therapy initiation on Day 3, and (ii) at the midpoint (Day 7 for cohort A and C, Day 10 for cohort B). Stool samples were collected on fecal occult blood test (FOBT) cards for all timepoints. Additional bulk stool scoop samples were collected at baseline for culturing work. Stool samples were stored at -80°C upon receipt at UCSF.

At the time of sequencing, there were 11 GO study participants who completed at least one cycle of treatment: 9 from Cohort A and 2 from Cohort C. 54 stool samples collected on FOBT cards from these 11 participants were used to extract DNA using Protocol Q from International Human Microbiome Standards (http://microbiome-standards.org). 300ng of normalized DNA was used in the Nextera DNA Flex library prep kit along with DNA UD Indexes Set A barcodes (Illumina) to assemble the metagenomic library. A blank FOBT card was included as a negative control. ZymoBIOMICS microbial community standard containing 8 bacteria and 2 yeasts (Zymo Research) was included as a positive control to assess bias and errors in the metagenomic library preparation. This mock community was significantly correlated with the theoretical composition (Pearson's correlation = 0.805; *p*-value 0.00496). The libraries were individually assessed with PicoGreen (ThermoFisher) and TapeStation 4200 (Agilent) for quantity and quality checks. The pooled library was sequenced using S1 flow cell on NovaSeq 6000 system (Illumina) at the Chan Zuckerberg Biohub.

**Selective culturing from colorectal cancer patient samples.** Baseline bulk stool scoop samples from 22 GO study participants were thawed on ice. Three participants did not submit proper baseline samples and were excluded. All processing and isolations were done under aerobic conditions. A single sample from each patient stool was taken and resuspended in $BHI^+$ media at a ratio of 15 ml/g stool. Stool samples were rigorously vortexed for 5 min and then allowed to settle for 10 min. Stool slurries were serially diluted 10-fold and and 1 ml samples of these various dilutions were spread on MacConkey agar on 150 x 15 mm petri plates. Agar plates were incubated aerobically at 37°C for 20 hours. The following day CFUs were counted and CFUs/gram of sample were calculated by dividing CFUs by dilution, and by sample weight. Isolated colonies were pooled by adding 1 ml of $BHI^+$ media supplemented with 15% (w/v) glycerol on top of the agar plate, resuspending and pooling colonies with a cell spreader, and recovering the cell suspension. A starter culture was initiated by using 50 µl of the cell suspension to inoculate 5 ml of anaerobically equilibrated $BHI^+$ media in Hungate tubes and incubated anaerobically at 37°C for 24 hours. The following day 50 µl of starter culture was used to initiate the 5-FU inactivation assay as described above.

**Data availability:** Sequencing data have been deposited under the NCBI BioProjects PRJNA576932 (RNA-seq, 16S rRNA gene sequences, and isolate genomes) and PRJNA720145 (GO Study metagenomes).

**Code availability:** Source code can be found at https://bitbucket.org/pbradz/preta.

**SUPPLEMENTARY TABLE LEGENDS**

**Supplementary Table 1. Minimal inhibitory concentration (MIC) data from a panel of human gut bacterial strains against fluoropyrimidines.**

**Supplementary Table 2. Generation and characterization of 5-FU resistant mutants.**

**Supplementary Table 3. *Escherichia coli* minimal inhibitory concentration (MIC) data.**

**Supplementary Table 4. RNA-seq metadata and sequencing statistics.**

**Supplementary Table 5. Differentially expressed bacterial transcripts in response to fluoropyrimidines.**

**Supplementary Table 6. Kinetic parameters for the activity of *E. coli* PreTA.**

**Supplementary Table 7. 16S rRNA gene sequencing metadata.**

**Supplementary Table 8. Prevalence and taxonomy of bacteria with a putative preTA operon across RefSeq bacterial genomes.**

**Supplementary Table 9. Prevalence and taxonomy of gut bacteria with a putative preTA operon from metagenome assembled genomes (MAGs).**

**Supplementary Table 10. List of primers used in this study.**

**Supplementary Table 11. List of *preTA* operons and sequences used for heterologous expression.**

**Supplementary Table 12. Gut Microbiome in Colorectal Cancer (GO) Study metadata (ClinicalTrials.gov Identifier: NCT04054908).**
